# Mitochondrial morphology and function in mitochondrial disease

**DOI:** 10.1101/2023.12.20.571319

**Authors:** Julie Faitg, Tracey Davey, Ross Laws, Conor Lawless, Helen Tuppen, Eric Fitton, Doug Turnbull, Amy E. Vincent

**Affiliations:** Wellcome Centre for Mitochondrial Research, Translational and Clinical Research, Faculty of Medical Sciences, Newcastle University, Newcastle, UK; Electron Microscopy Research Services, Newcastle University, Newcastle, UK; John Walton Muscular Dystrophy Research Centre, Translational and Clinical Research, Faculty of Medical Sciences Newcastle University, Newcastle, UK; NIHR Biomedical Research Centre Research Centre, Translational and Clinical Research, Faculty of Medical Sciences Newcastle University, Newcastle, UK

**Keywords:** Mitochondria, shape, morphology, muscle pathology, mitochondrial disease

## Abstract

Mitochondria play a crucial role in maintaining cellular health. It is interesting that the shape of mitochondria can vary depending on the type of cell, mitochondrial function, and other cellular conditions. The morphology of this network is maintained by the balance between mitochondrial fusion and fission, necessary for the proper functioning of the mitochondrial energetics as well as the maintenance of mitochondria DNA (mtDNA) (Chan, 2006; Friedman and Nunnari, 2014; Glancy et al., 2015). If the balance between the two processes is slightly perturbed, dramatic changes in mitochondrial morphology can occur.

However, there are limited studies that link functional assessment with mitochondrial morphology evaluation at high magnification, even fewer that do so *in situ* and none in human muscle biopsies. Therefore, we have developed a method which combines functional assessment of mitochondria through Cytochrome c Oxidase (COX) histochemistry, with a 3D electron microscopy (EM) technique, serial block-face scanning electron microscopy (SBFSEM). Using this technique, we examine the relationship between 3D mitochondrial morphology and mitochondrial activity in human skeletal muscle.

Here we apply COX-SBFSEM to muscle samples from patients with single, large-scale mtDNA deletions, a cause of mitochondrial disease. These deletions cause oxidative phosphorylation deficiency, which can be observed through changes in COX activity, and a typical mosaic pattern of enzyme deficiency with both COX normal and deficient fibres present in the same biopsy.

Using COX-SBFSEM, we can distinguish three distinct mitochondrial populations within muscle fibres: COX normal (+), COX intermediate (±) and COX deficient (-) mitochondria. COX+ mitochondria were larger and more complex than the COX- mitochondria, However, in fibres that were generally COX-, there was no apparent difference between the three types of mitochondria and all mitochondria in fibres that are COX- overall are fragmented and spherical. One of the main advantages of combining 3D-EM with the COX reaction is the ability to look at how per-mitochondrion oxidative phosphorylation status is spatially distributed within muscle fibres. Here we show a robust spatial pattern in fibres that are generally COX+ and that the spatial pattern is less clear in fibres that are predominantly COX± and COX- .

## Introduction

In skeletal muscles there are two mitochondrial populations, subsarcolemmal mitochondria (SS), located beneath the cell membrane, and intermyofibrillar mitochondria (IMF), located between myofibrils (Ogata and Yamasaki 1985). SS mitochondria are generally more spherical, with some exceptions of elongated mitochondria which protrude into the intermyofibrillar space (Picard, White and Turnbull 2013, Vincent, White et al. 2019). IMF mitochondria, on the other hand, appear to be organised into an interconnected reticulum, with more complex mitochondria (Picard, Hepple and Burelle 2012, Vincent, White et al. 2019). It is also worth noting that, as well as morphological differences, these two populations exhibit functional specialisation including differences in capacity to generate ATP (Feirrera et al., 2010).

Mitochondrial DNA (mtDNA) mutations are associated with mitochondrial oxidative phosphorylation (OXPHOS) dysfunction in skeletal muscle. These mtDNA mutations can be inherited, occur sporadically early in embryogenesis or arise as a result of nuclear gene mutations that impact mtDNA maintenance. Since mtDNA is present in hundreds to thousands of copies per cell, a mutation in one or a small number of mtDNA copies has negligible impact. However, when present in higher proportions of the mtDNA within a single cell, a mtDNA mutation may exceed a biochemical threshold (Rossignol, Faustin et al. 2003) and cause mitochondrial dysfunction. MtDNA contains genes encoding subunits of complex I (CI), complex III (CIII), complex IV (CIV) and complex V (CV), as well as tRNAs and rRNAs for mitochondrial translation. As such, mtDNA mutations may give rise to deficiency in CI, III, IV or V depending on which genes are impacted, with cytochrome c oxidase (COX) or CIV being the most commonly observed deficiency alongside CI deficiency (DiMauro, Zeviani et al. 1985, Morgan-Hughes 1986, Nonaka, Koga et al. 1989).

In patients with mitochondrial myopathy COX activity is heterogeneous between cells, with a mosaic pattern of enzyme deficiency where both COX normal and deficient fibres are present in the same biopsy, and this is easy to observe using histology. Mitochondrial dysfunction exhibits a segmental pattern within muscle fibres, with dysfunctional mitochondria surrounded by healthy mitochondria, both longitudinally and in the transverse orientation (Dimauro, Nicholson et al. 1983). Furthermore, mtDNA mutations can impair Ca^2+^ handling (Eisner, Caldwell et al. 2017), mitochondrial dynamics (Ranieri, Brajkovic et al. 2013) and patients with mtDNA mutations are found to have altered mitochondrial morphology and cristae organisation in skeletal muscle biopsies (Vincent, Ng et al. 2016, Vincent, White et al. 2019).

In a recent study by Vincent, White et al. (2019), mitochondrial morphology and network organization in human skeletal muscle were quantitatively analysed. Mitochondrial disease patients with mtDNA mutations were found to have an elevated frequency of simple mitochondria and mitochondrial nanotunnels (Vincent, White et al. 2019). However, the limitation of this work was that OXPHOS deficiency could not be detected, so any link between mitochondrial morphology and function could not be observed. Thus, it is unclear how mitochondrial morphology changes as OXPHOS deficiency progresses from normal to intermediate to deficient to ragged red fibres (RRF) and mainly what is happening in 3D space regarding the spread of dysfunction.

The successful development of the COX-SBFSEM technique on mouse tissue (Faitg, Davey et al. 2020) now allows us to apply this to patient tissue and correlate mitochondrial function and morphology *in situ*. This study aimed to extend the work done in SBF-SEM and use the COX-SBFSEM technique to compare OXPHOS deficient muscle fibres with normal muscle fibres in mitochondrial disease patients.

## Methods

### Cohort clinical characteristics

Quadriceps muscle samples were collected through the AIMM Trial (AIMM Trial Group, *Trials* (2022); REC ref: 18/NI/0199, ISRCTN: 12895613). Baseline samples (∼1mg) from four patients with single, large-scale mtDNA deletions (Table 1) were processed for EM. Mutation load was determined as part of the AIMM Trial. Deletion breakpoints were confirmed from muscle DNA requested from the Newcastle Mitochondrial Research Biobank (NMRB050; REC ref: 21/NE/0204). Figure 1 shows schematically which genes are deleted in each patient.

**Figure 1.**
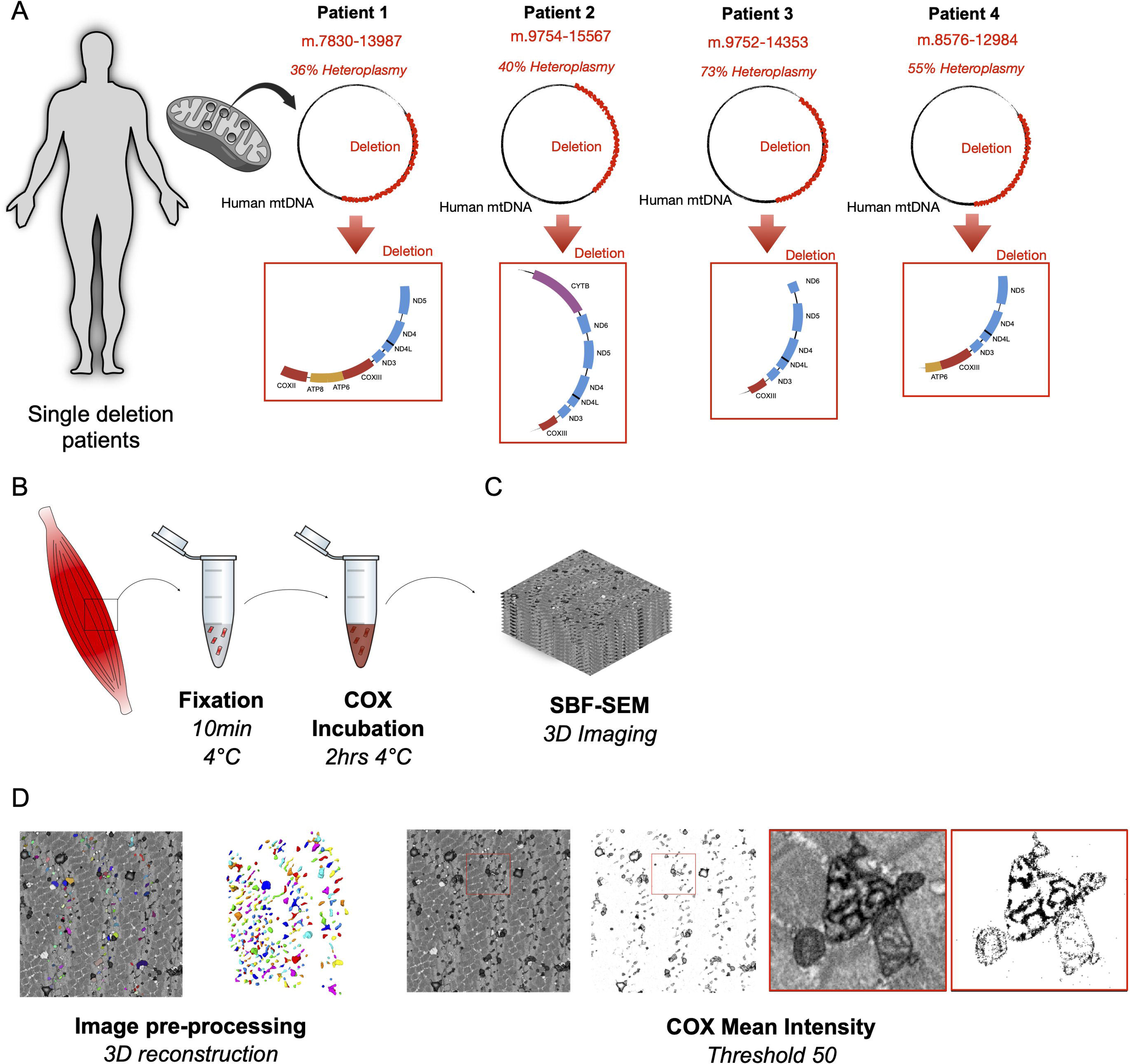
Schematic showing work flow for experiments. **(A)** Schematic of study design involving single deletion patients. **(B)** Muscle biopsy and teasing fibre. fixation and imaging of bundles muscle fibre and mitochondria in transversal orientation. **(C)** Z stack at EM resolution from serial block face scanning electron microscopy (SBF-SEM) used for 3D reconstructions (dimensions of 20 µm x 20 µm). **(D)** Image processing and 3D reconstruction of individual mitochondria (left) and thresholding for COX intensity including high resolution image of COX positive and negative mitochondria (left).

**Table 1.** Clinical and genetic information for subjects included in this study.

| **Patients** | **Sex** | **Age at biopsy** | **Genetic defect** | **Mutation load** |
| --- | --- | --- | --- | --- |
| **P1** | F | 47y | Deletion size ~ 6.1kb<br>Breakpoints: 7830 - 13987 | 36% |
| **P2** | F | 43y | Deletion size ~ 5.8 kb<br>Breakpoints: 9754 - 15567 | 40% |
| **P3** | M | 26y | Deletion size: ~4.6kb<br>Breakpoints: 9752-14353 | 73% |
| **P4** | F | 49y | Deletion size ~ 4.5kb<br>Breakpoints: 8576-12984 | 55% |

f, female; m, male; y, years; Kb, Kilobases.

### COX - SBFSEM processing and imaging

Human muscle samples were processed with the COX-SBFSEM as previously described (Faitg, Davey et al. 2020). Briefly, a short fixation for 10min at 4°C, followed by incubation with COX medium for 2 hours at 37°C in the dark. Samples were fixed for 10min at 4°C and washed three times with Sorenson’s buffer in the microwave (3x 150W 40s per step as described by Faitg, Davey et al. 2020). The samples were incubated with the COX reaction for 2 hours at 37°C in the dark (Faitg, Davey et al. 2020). The tissue was processed using a heavy metal protocol. Briefly, samples were subsequently fixed, and membranes were stained with 3% potassium ferrocyanide and 2% osmium tetroxide for 1 hour at room temperature (RT). Following osmium fixation, the remainder of the phosphate buffer was removed by washing in 0.1M Sorenson’s buffer (3 x 40 sec per step, 150W, (Faitg, Davey et al. 2020)). The samples were placed into a contrast enhancer 0.1% thiocarbohydrazide (TCH) filtered at RT for 20 min and immersed in 2% osmium tetroxide for 30 min at RT. Finally, samples were placed into 1% uranyl acetate overnight at 4 °C. On day two, all samples were washed in several changes of distilled water (ddH_2_O; 5 x 3min RT and microwave 3x40s 150W, (Faitg, Davey et al. 2020)). Following. Following these washes, immersion for 30 min at 60 °C in lead aspartate solution (previously heated for 30 minutes: 0.12g of lead nitrate in 20mL aspartic acid) (Wilke, Antonios et al. 2013) was carried out. Dehydration was performed in a graded series of acetone from 25% to 100% and impregnation in increasing concentrations of Taab 812 hard resin (25% to 100%) in acetone with several changes to 100% resin. Samples were embedded in 100% fresh resin and left to polymerise at 60°C for a minimum of 36 hours (Figure 1B). After polymerisation, the resin blocks were trimmed to approximately 0.75mm × 0.75mm and sectioned as described by Faitg, Davey et al. (2020) to identify the regions of interest (ROIs) before performing SBFSEM (Faitg, Davey et al. 2020). For each sample, transversely orientated muscle fibres were visually selected to have as many COX-normal and COX-deficient fibres in close proximity to each other as possible (i.e., minimise re-localisation and imaging time). One ROI for IMF mitochondria was selected for each fibre and imaged. The samples were sectioned and captured in a series of images (400 images per stack) at 70nm sectioning thickness. The image resolution achieved with SBF- SEM was 2000 x 2000 pixels and a pixel size of 0.07μm x 0.07μm in the x,y dimensions (Faitg, Davey et al. 2020).

### Image analysis pre-processing and mitochondrial 3D reconstruction

Image stacks from Digital Micrograph (Gatan) were normalised as described in Faitg et al. 2020 and converted to Tiff files. Briefly a standard operation, which consists of 3 steps: (i) obtain the mean and standard value of intensities for the whole dataset, (ii) for each image, and finally (iii) shift/stretch each image such that its mean/standard values matched the mean/standard of the whole dataset (Faitg, Davey et al. 2020). Once in the right format, the mitochondrial segmentation was completed throughout the stack in all three dimensions, across two sarcomeres from A-band to A-band, to generate 3D reconstructions (MIB, Helsinki version 2.702 (Belevich, Joensuu et al. 2016)). The mitochondria were manually traced using the ‘brush’ tool in MIB software (Belevich, Joensuu et al. 2016). After which, the mitochondria were reconstructed in 3D to extract volume and surface area data with Amira (AMIRA image analysis software, version 2020.3). Surface area and volume for each mitochondrion was extracted using AMIRA. Mitochondrial Complexity Index (MCI): [MCI=((SA^1.5^)/4πV))^2^] a 3D equivalent to form factor which quantifies mitochondrial branching, was calculated as described previously by (Vincent, White et al. 2019).

### Analysis of COX labelling intensity

The electron microscopy 3D images stacks were normalised using MIB as mentioned above. For measurements of COX specific activity, the threshold was manually set to include only COX specific enzyme cytochemical product. An intensity threshold of 50 was chosen so that only the COX specific precipitate and therefore only COX positive cristae were visible (Figure 1D). The mean intensity per pixel was obtained for each individual mitochondrion from all fibres. The mean intensity value was close to 0 for black and 255 for white.

### Statistical analysis of mitochondrial morphology and COX-EM data

A normality test was performed on all datasets for the four patients and histograms plotted in Prism v9.0 (Graph Pad) to assess normality. The data were not normally distributed, precluding the use of parametric statistical tests. Therefore, for multiple comparisons (more than two groups) the Kruskal-Wallis test or Mann-Whitney (for two groups only) were applied, to examine the main effects, followed by posthoc tests using the two-stage step up method of Benjamini, Krieger, and Yekutieli to correct for multiple comparisons (17p < 0.05, q < 0.05).

### Multivariate analysis and machine learning

To describe the observed distributions of mitochondrial morphology from the four patients, partial least-squares discriminant analysis (PLS-DA) and clustering analysis using a hierarchical ward algorithm presented as a heatmap were generated using the MetaboAnalyst 5.0 (Pang et al., 2021). The variable importance in projection (VIP) scores were extracted for each mitochondrial feature for the PLS-DA models.

### Neighbourhood analysis

To visualise whether deficient mitochondria cluster together, the x,y,z coordinates of every single mitochondrion and their respective COX activity were put together in the same 3D scatter plot. The source code can be found in supplemental Appendix 1.

## Results

### COX activity at a single mitochondrion level

The fibres were selected by eye during the imaging step. COX normal fibres were easily recognisable by the dark, dense precipitate within the mitochondrial cristae, in comparison COX deficient fibres lacked this precipitate (Figure 2A). The spectrum of COX intensity of individual mitochondria was analysed for each fibre (Figure S1). An intensity value of 0 indicates black or electron dense, corresponding to a high concentration of COX and 255 indicates white or absence of the COX enzyme. For all patients, three groups of fibres grouped by their mean COX intensity are visible (Figure 2B-F). The comparative COX intensity distributions for all patients are shown in Figure 2B, highlighting the heterogeneity in distribution between patients.

**Figure 2.**
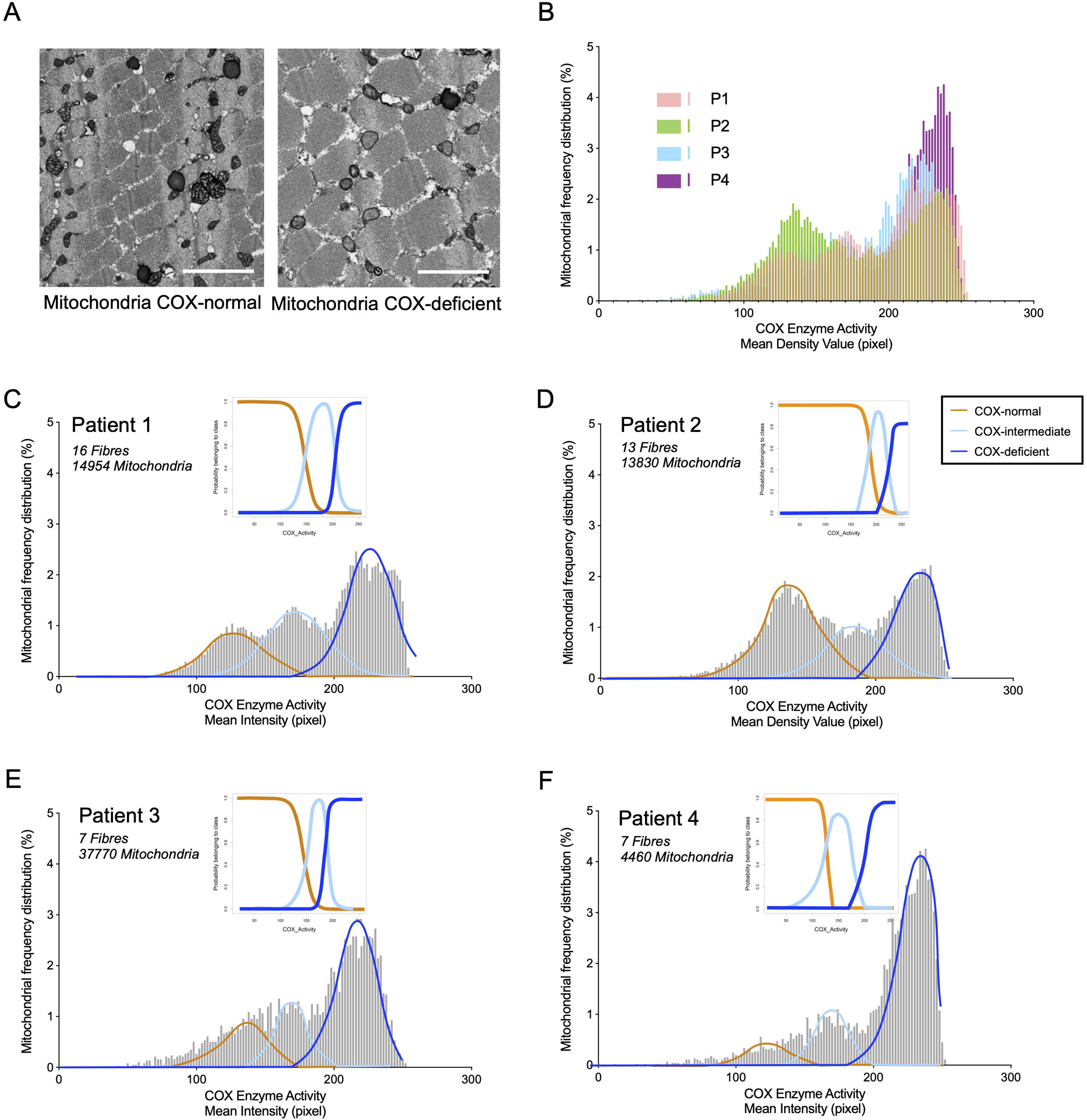
Mitochondrial COX activity falls into three distinct groups. **(A)** Single SBF-SEM image of a COX normal fibre (left) and a COX-deficient fibre (right). **(B)** Frequency distribution of mitochondrial COX activity from individual fibres of all single, large-scale mtDNA deletion patients. Patients, n=4; Fibre, n=X; Mitochondria, n=-xxx. **(B-E)** Gaussian Mixture Model to P1(B), P2(C), P3(D), P4(E) data, each mitochondrion belongs to the orange, light blue, or blue cluster fit to COX activity distributions from Patient 1: Fibres n=16, 14954 mitochondria. Patient 2: Fibres n=13, 13830 mitochondria. Patient 3: Fibres n= 7, 3770 mitochondria. Patient 4: Fibres n=7, 4460 mitochondria.

Using Gaussian mixture modelling, an unsupervised clustering method, based solely on COX intensity, the mitochondria split into three clusters (orange, light blue, or blue in Figure 2C- F). A probability of each mitochondrion belonging to the orange, light blue, or blue cluster was calculated, the intersection between these curves represents the mitochondria where there is some uncertainty in classification and so a proportion of mitochondria cannot be classified (Figure 2C-F). In order, to not lose large numbers of mitochondria, the intersection between the orange and light blue lines was used as a threshold between COX-normal and COX-intermediate mitochondria and the intersection between the light-blue and blue lines as the threshold between COX-intermediate and COX-deficient mitochondria, thus allowing us to classify all mitochondria as either COX-normal, -intermediate or -deficient.

Interestingly, the threshold for P2,3&4 for COX deficiency is higher than for any of the other patients. This is likely due to this patient harbouring a deletion involving more CI genes and fewer CIV genes. P2,3&4 would have been classified as a class II deletion based on work by Rocha et al (2018). Importantly, this patient will have some fibres and some mitochondria that will have only CI deficiency and appear COX-normal in this assay. In comparison P1 have a deletion that would be classified as a class I deletion and we would expect mainly cells with CI and CIV deficiency (Rocha et al., 2018), therefore the majority of OXPHOS deficient cells and mitochondria should be picked up as COX deficient by the COX-SBFSEM assay.

It is interesting to note that as expected, when we compared the proportion of COX normal, intermediate, and deficient mitochondria in each fibre there is a distinct pattern (COX- normal fibres include mostly COX normal mitochondria) in the fibres classified as COX positive or deficient by eye (Figure 3). The fibre classifications were based on the percentage class of the individual mitochondria in a fibre (Figure 3 and Table S1).

**Figure 3.**
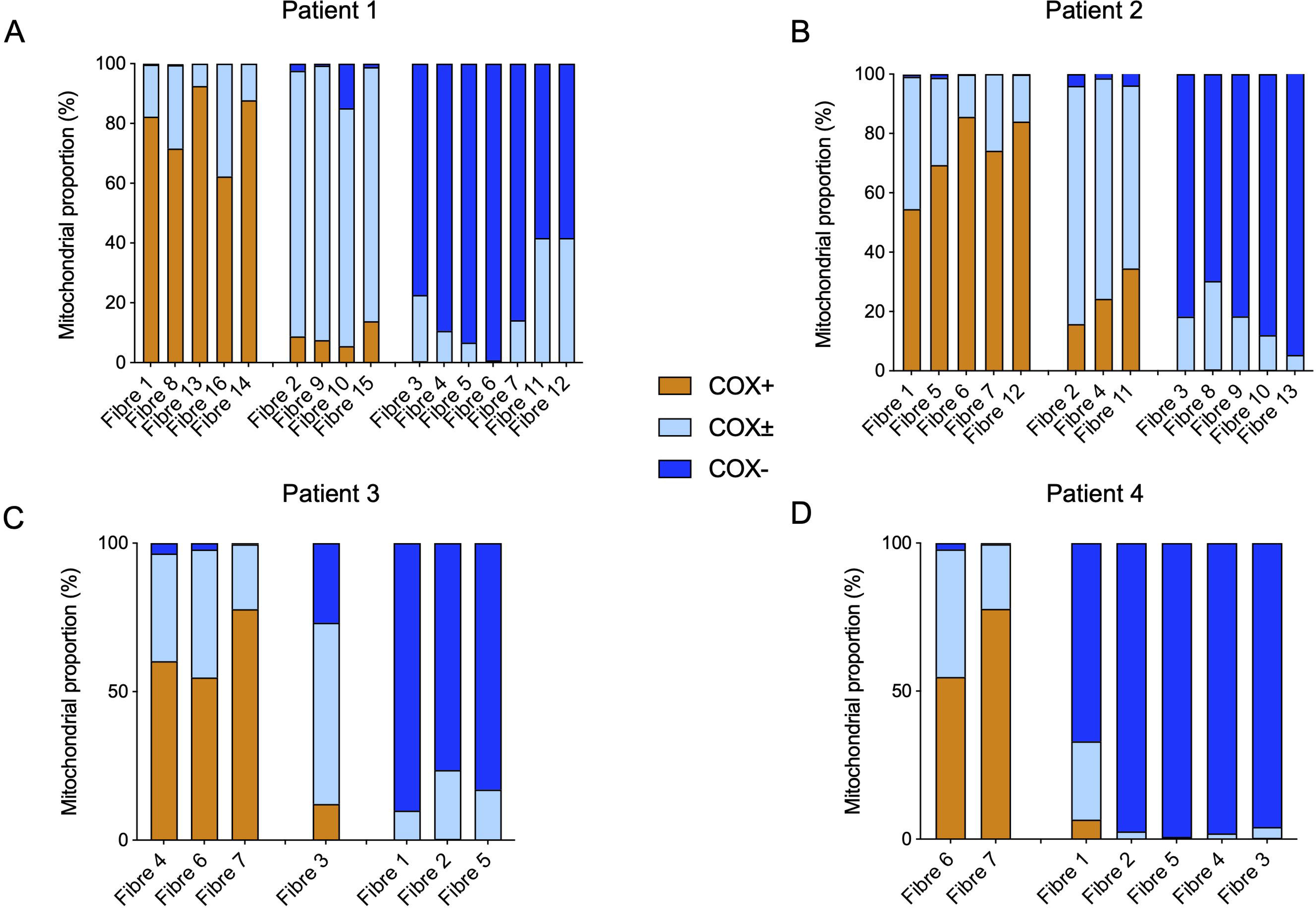
Proportion of mitochondria classified as COX positive, intermediate or deficient in each fibre. Proportion of Mitochondria classified as COX positive, intermediate and deficient proportion within each fibres from (**A**) patient 1, (**B**) patient 2, (**C**) patient 3 and (**D**) patient 4. COX positive mitochondria are represented in orange; COX intermediate mitochondria are represented in light blue; COX negative mitochondria are represented in blue. Typically fibres exhibiting more than 50% COX positive mitochondria were visually classified as COX positive fibres, those exhibiting more than 50% COX intermediate mitochondria were visually classified as intermediate fibres and those exhibiting more than 50% COX deficient mitochondria were visually classified as COX deficient mitochondria.

### The relationship between COX activity and mitochondrial morphology within muscle fibres

Since previous work in cell models has found links between mitochondrial morphology and function, we sought to compare mitochondrial morphology between COX-normal, COX- intermediate and COX-deficient fibres across the four patients. The mitochondria from COX- normal fibres showed a large spectrum of volume and complexity in the four patients and between individual fibres (**^⍰^**p<0.05**, ^⍰⍰⍰⍰^**p<0.0001, Figure 4A-C). Concerning the COX- deficient fibres, mitochondrial volume was significantly smaller in P1-3 but greater in P4 (Figure 4A and B). No fibre exhibited a mean mitochondrial volume greater than 0.05 µm^3^ excepted for P4 (Figure 4A).

**Figure 4.**
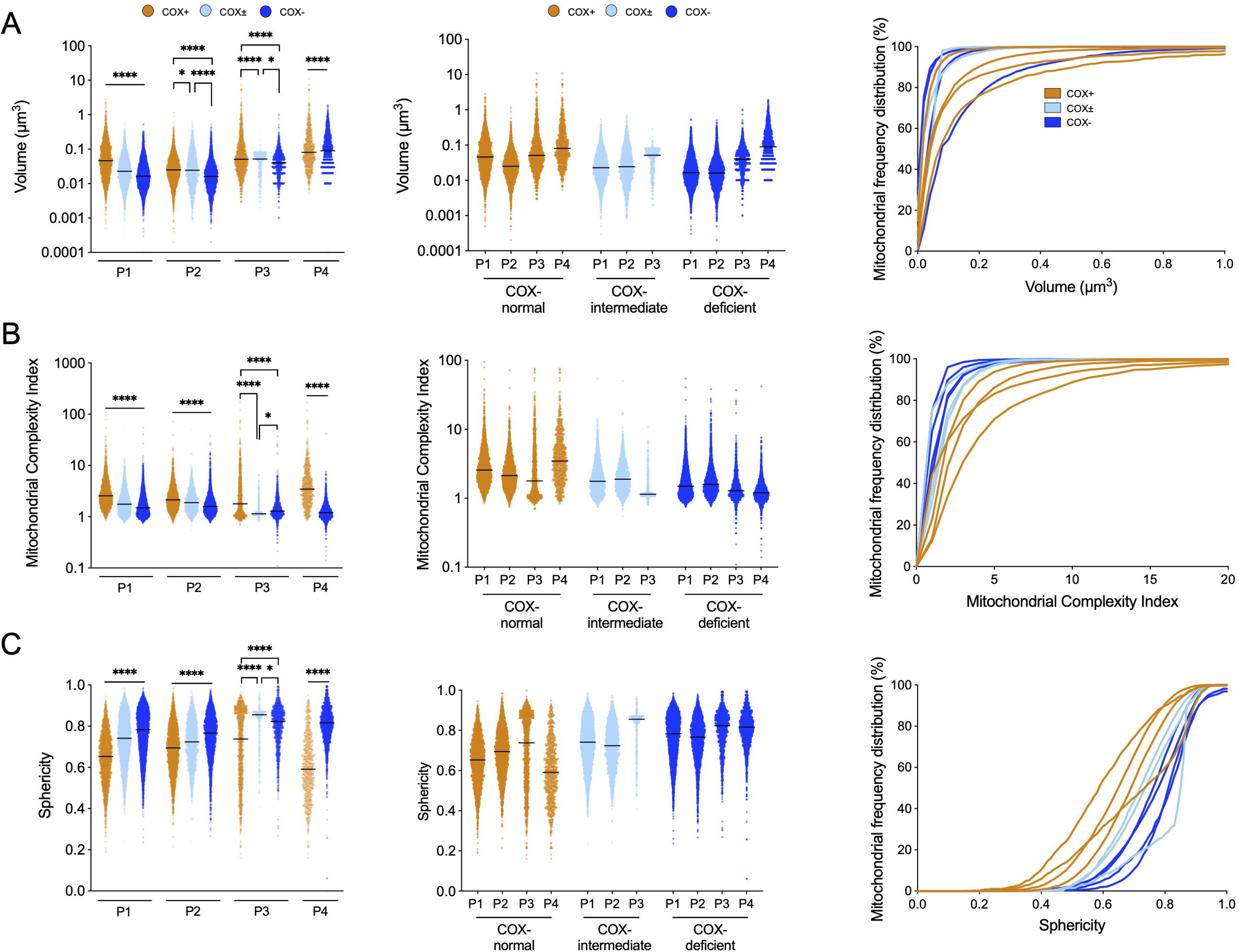
General morphological comparison between mitochondria of COX-normal, intermediate, and deficient fibres from all patients. **(A-C)** Median mitochondrial volume , MCI **(B),** and sphericity (C) with their respective cumulative frequency distribution of the COX­normal, COX-intermediate, and COX-deficient fibres. Mitochondrial complexity Index (MCI) is a three dimensional metric of mitochondrial shape complexity. MCI is an analogous to sphericity and scales with mitochondrial shape complexity, including branches and increased surface area relative to volume. Data are presented as median with 95% Cl. Kruskal-Wallis test followed by post-hoc tests using the two-stage step-up method of Benjamini, Krieger, and Yekutieli to correct multiple comparisons (p < 0.05, q < 0.05).

Based on MCI, mitochondria were significantly simpler in COX deficient fibres compared to COX-normal and -intermediate, with the exception of COX-intermediate fibres in P3 which were slightly simpler (Figure 4B). Interestingly, all COX-deficient fibres for each patient showed average mitochondrial sphericity greater than 0.6 (Figure 4C).

We next wished to look at the relationship between the COX activity values for each fibre and the MCI values for each fibre. We did this by plotting the mean and SEM for each fibre for COX activity against MCI (Figure 5). In patient 1 it appears that MCI tends to increase as the COX activity increases (value decreases) (Figure 5A). In patient 2 this is also true however the range of MCI values is relatively narrow and so the relationship does not appear as strong (Figure 5B). In patient 3 we see the data distributed in an L shape with a cluster of fibres with similar MCI and a range of COX activity and then a few fibres with higher COX activity and a range of MCI values which are higher than for the other group of fibres (Figure 5C). In patient 4 we can also see the pattern of MCI increasing with COX activity (Figure 5D).

**Figure 5.**
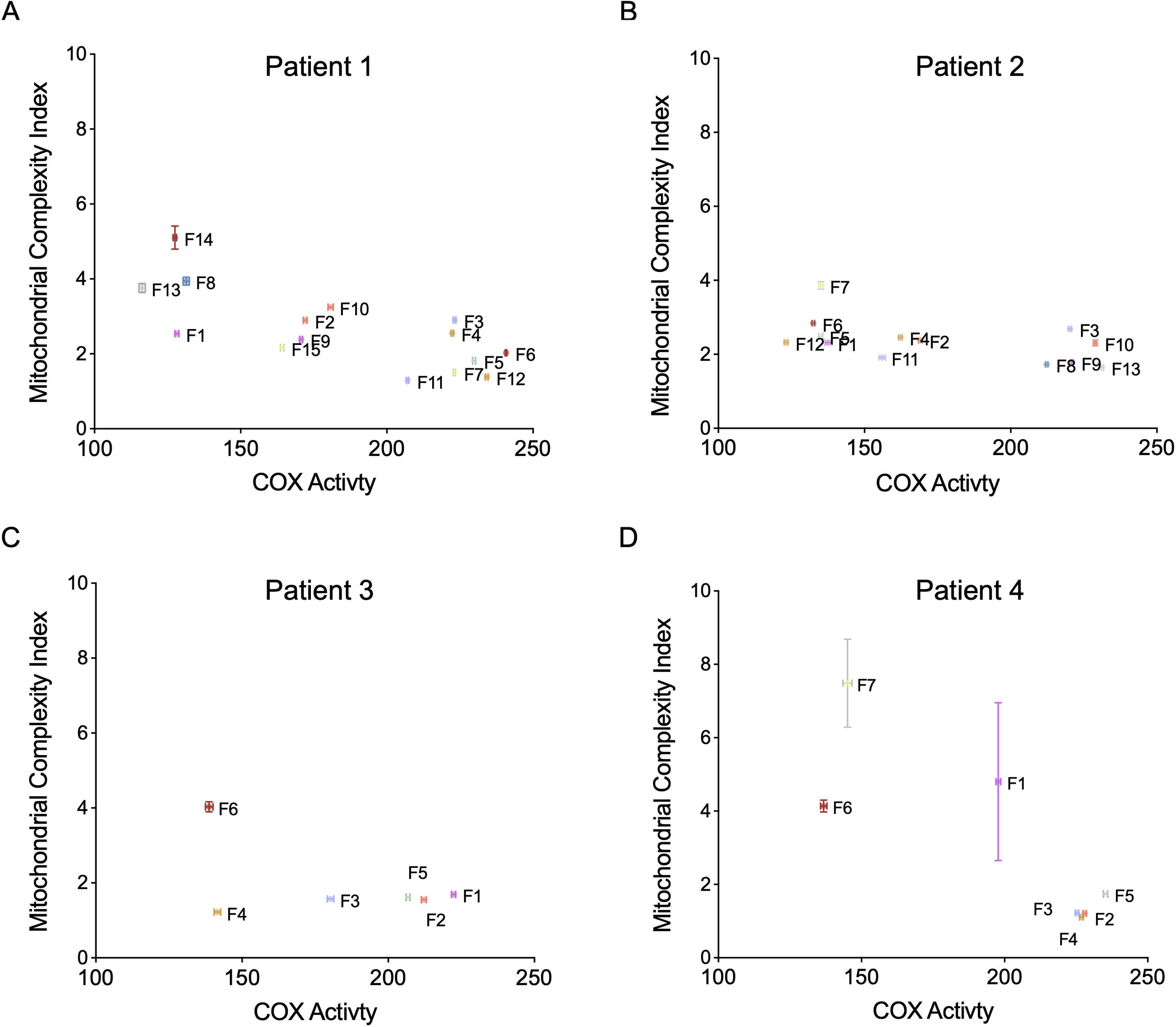
Effect of Single deletion mutation on mitochondrial morphology and function for each patient. **(A-D)** Mitotypes illustrating the difference in COX activity and MCI between fibres (mean ± SEM), for patient 1 (A), patient 2 (8), patient 3 (C) and patient 4 (D).

We sought to determine whether mitochondrial morphological parameters can be used to differentiate COX-normal from COX-deficient fibres in patients with mitochondrial disease, as they were shown to differentiate Controls and Patients by Vincent, White et al. (2019). To do this we applied multivariate analysis methods similar to those used by Vincent, White et al. (2019). Using PLS-DA the model captured 92.1% of the total variance in the dataset (Figure 6A). The COX-normal fibres were more widely distributed, with the COX-deficient fibres mostly contained within the red ellipse, with the exception of 3 points falling outside (Figure 6A). For each parameter, the rank ordering of the VIP scores (a score > 1 is considered significant) identified the most important morphological features that differentiate between COX-normal and COX-deficient fibres. (Figure 6B). The top four features for all patients together when comparing COX positive vs negative fibres, were (1) the MCI mean, (2) the MCI median, (3) the volume mean and (4) the sphericity median (Figure 6B), where the first 3 had highest values for COX normal fibres. The top features were also generated for each patient separately, similarly, to compare COX normal vs deficient fibres (Figure S2). A heat map was generated based on group means of each fibre group, COX-deficient fibres had a higher sphericity compared to the COX-normal and other parameters were all lower for all patients combined (Figure 6C) and for each patient individually. This demonstrates for the first time that a high proportion of morphologically simple and spherical mitochondria represent a signature of mitochondrial OXPHOS deficiency in human muscle fibres.

**Figure 6.**
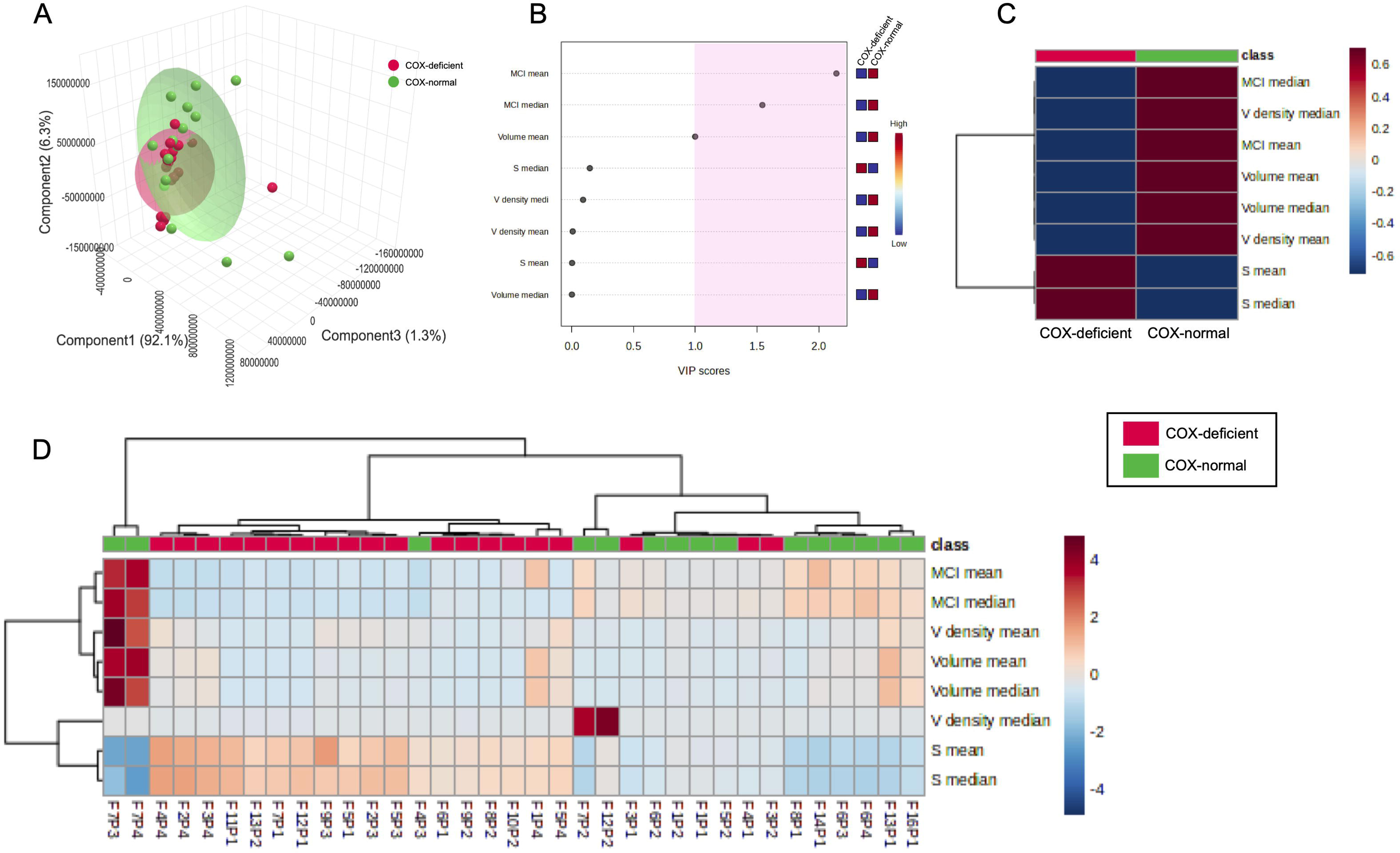
Multivariate analysis of mitochondrial morphology between individual normal and deficient fibres from all patients. **(A)** Partial Least Squares Discriminant Analysis (PLS-DA) highlights morphological variables specific to the differentiation of COX­ normal and COX-deficient fibres. On the left, the PLS-DA score plots were presented for each type of variable. The model explains 92.1% (PC1) + 6.3% (PC2) + 1.3% (PC3) = 99.7% of the variance. **(B)** The variable importance in projection or VIP score for the morphology parameters used in the PLS-DA regrouping the 4 patients together. The parameters with VIP score > 1 are considered significant. The coloured boxes on the right indicate the relative concentrations of the corresponding metabolite in each group under the current study. **(C-D)** Heatmap across group average **(D),** and all subjects and parameters **(C),** with dendrograms illustrating hierarchical clustering of pattern similarity across morphological parameters and samples (top) (Euclidean distance measure, Ward clustering algorithm). The colours indicate the relative quantitative value, where red indicates a higher value, and blue indicates a lower value. The COX-normal fibres: <u>Patient 1</u>: n= 5; <u>Patient 2:</u> n=5; Patient 3: n=3; Patient 4: n=2. The COX-deficient fibres: <u>Patient 1</u>: n= 7; <u>Patient 2:</u> n=5; Patient 3: n=3; Patient 4: n=5.

Hierarchical clustering highlights the similarities and differences between COX-normal and COX-deficient fibres (Figure 6D). This unsupervised analysis which clusters fibres based on similarities between samples and functional parameters shows that COX-normal vs COX- deficient fibres segregate into two distinct clusters. One exception came from three COX- deficient fibres, which had lower mitochondrial sphericity (Fibre 3 P1, Fibre 4 P1, Fibre3 P2) than the other deficient fibres (Figure 6D).

### The relationship between COX activity and mitochondrial morphology in individual mitochondria

We initially compared the morphology of COX-normal, intermediate, and deficient mitochondria within single normal, intermediate and deficient fibres (Figure 7A-C). Within normal fibres from all patients, the COX-normal mitochondria were on average significantly larger and more complex than COX-intermediate mitochondria, which were in turn larger than COX-negative mitochondria (Figure 7A-C). Regarding the intermediate fibres, the COX- intermediate mitochondria were larger and more complex, while the COX-deficient mitochondria were smaller and more spherical within these fibres. In addition, within The deficient fibres, mitochondrial COX-deficient were also smaller and more spherical than the others. However, when looking at individual patients, no significant differences were seen in MCI between these 3 groups for individual patient, however, the N was too small to draw a robust conclusion (Figures S3-S6).

**Figure 7.**
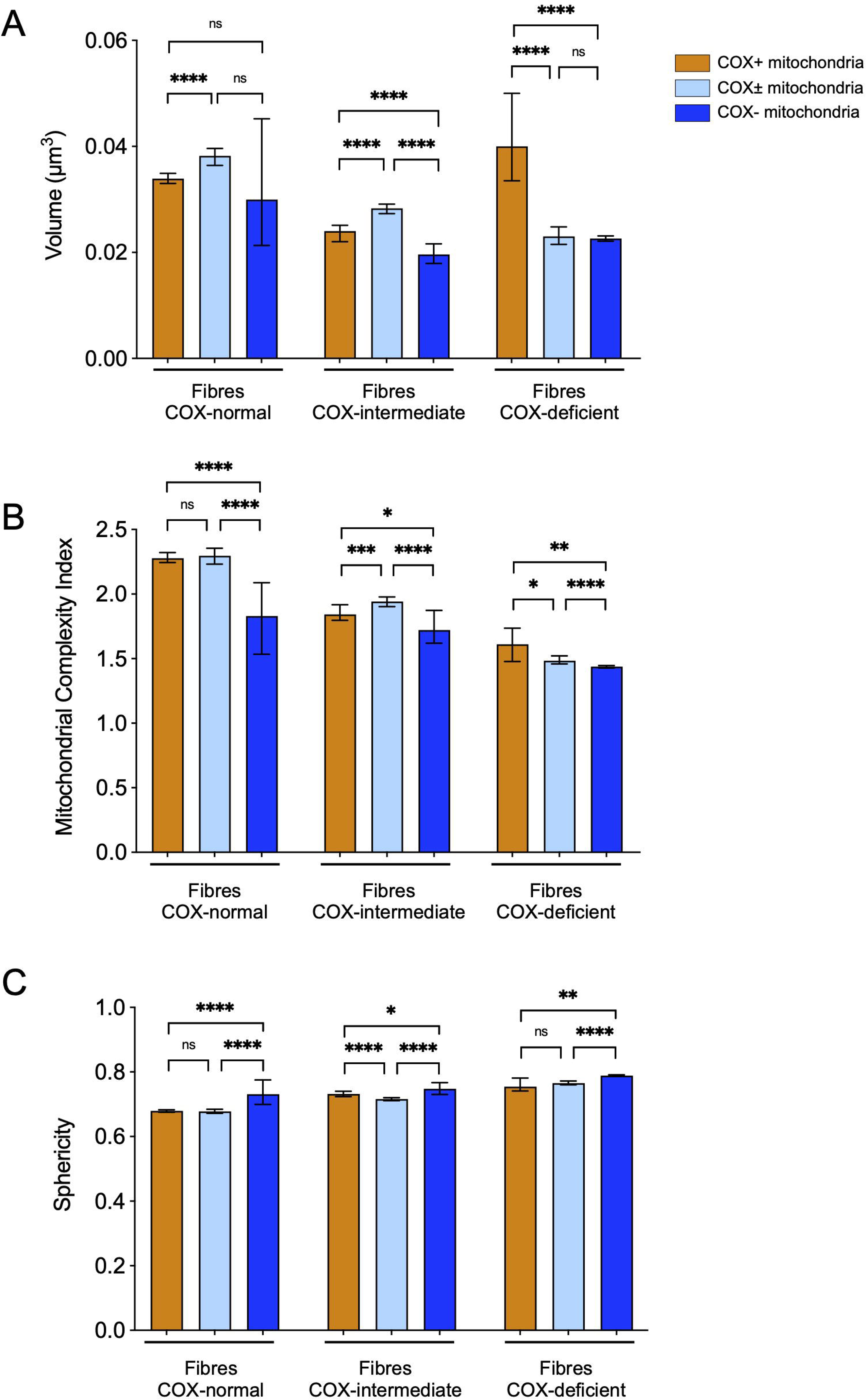
Morphological comparison of the three distinct mitochondria groups present within each fibre. This involve analysis of the COX-normal, COX-intermediate, and COX-deficient fibres of all the regrouped patients in order to draw a conclusive comparison. **(A-C)** Median mitochondrial volume , MCI **(B),** and sphericity **(C)** with their respective cumulative frequency distribution of the COX­normal, COX-intermediate, and COX-deficient fibres. Data are presented as median with 95% Cl. Kruskal-Wallis test followed by post-hoc tests using the two-stage step-up method of Benjamini, Krieger, and Yekutieli to correct multiple comparisons (p < 0.05, q < 0.05).*p<0.05,**p<0.005, *** p<0.0005, ****p<0.0001

### COX deficiency and mitochondrial morphology with spatial resolution

The intracellular COX heterogeneity distinguished above is illustrated in Figure 8. Within the COX-normal fibre, COX-normal and intermediate mitochondria are grouped spatially in a strong pattern (Figure 8, Figure S7). This pattern can be also seen in the COX-intermediate fibre. Interestingly, most of the COX-deficient fibres exhibited a mix of COX-deficient and intermediate mitochondria, without a spatial pattern. The distribution tends to confirm the spread of the disfunction across the fibre, this is also surprising as it suggests that mitochondria are very separate and are becoming deficient independently. Much like the mosaic pattern at the tissue level (Figure 8, Figure S7).

**Figure 8.**
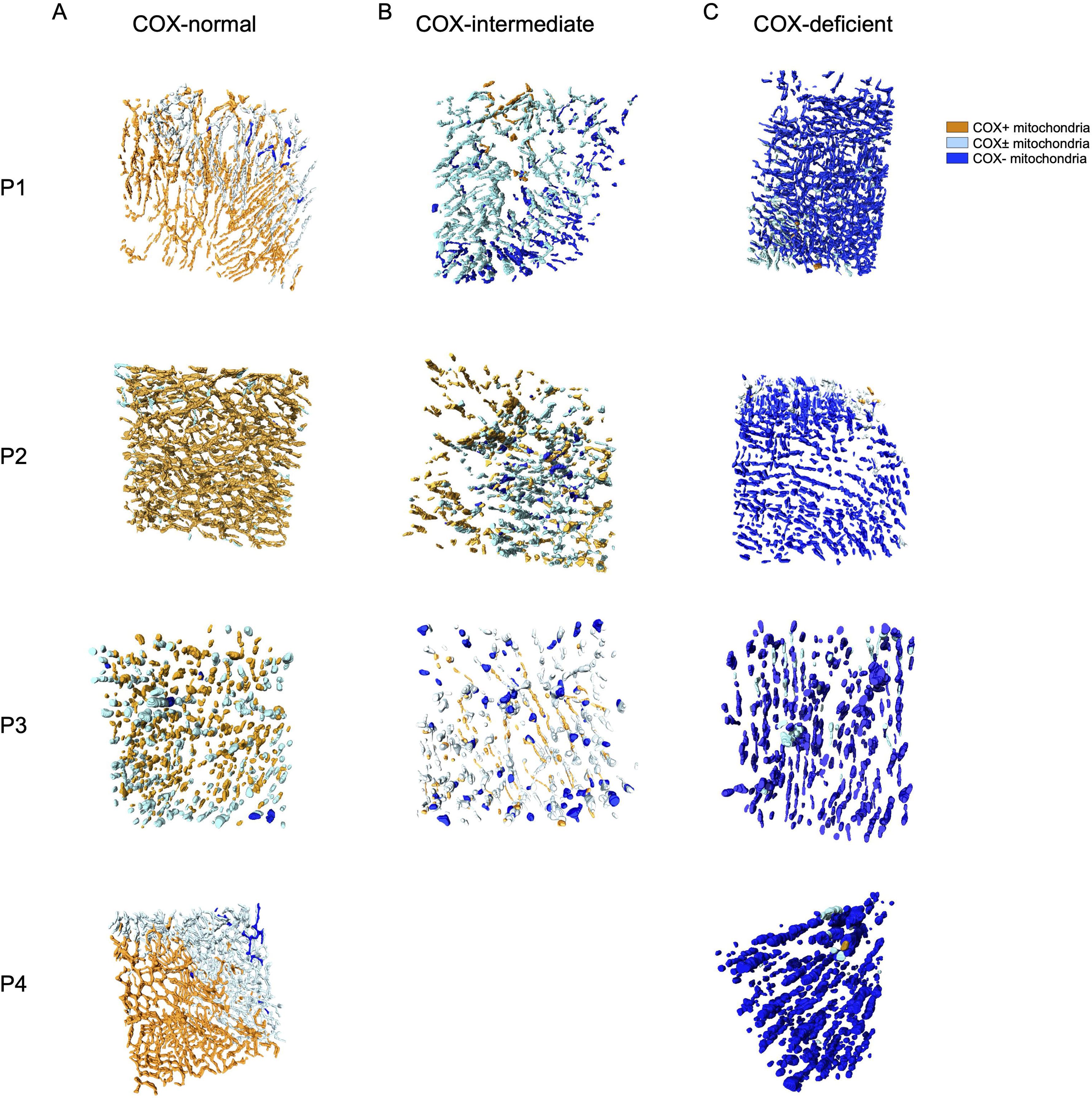
COX activity and spatial distribution of the mitochondrial COX activity and their respective morphologies within a COX-normal/COX-intermediate and COX-deficient fibres from Patient 1,2,3&4. The 3D reconstruction figures of a COX-normal (Figure ?A), COX-intermediate (Figure 7B) and COX-deficient (Figure 7C) fibre demonstrate coloured mitochondria according to their COX activity. 3D reconstruction of mitochondrial COX activity across two sarcomeres in COX normal (A), COX-intermediate (B) and COX-deficient (C) fibres demonstrate coloured mitochondria according to their COX activity. Mitochondria are colour coded dependent on COX activity as indicated by the scale Cluster of COX-normal and intermediate mitochondria can be distinguished by the dotted circle.

When we look at mitochondrial morphology using MCI in a similar way, we find very little pattern spatially in COX-normal, COX-intermediate and COX-deficient fibres (Figure 9). However, to better visualise the morphology and COX activity together we generated videos of the fibres in Figures 8 and 9 where the 3D reconstruction has been color-coded based on COX activity (COX-normal: orange, COX-intermediate: light blue, COX-deficient: dark blue). In Video 1 you can see a COX normal fibre from patient 1 which has highly branched normal mitochondria with a small number of more fragmented deficient mitochondria, with a strong spatial pattern to mitochondrial function. In Video 2 we present the intermediate fibre from patient 4, which shows a strong spatial pattern regarding their activities and a mix of mitochondrial morphologies, where the COX deficient mitochondria still exhibit simple mitochondrial morphology. Finally, in Video 3 we present a COX deficient fibre from patient 2, where most mitochondria are deficient and relatively simple in morphology but a strong spatial pattern in COX activity is visible.

**Figure 9.**
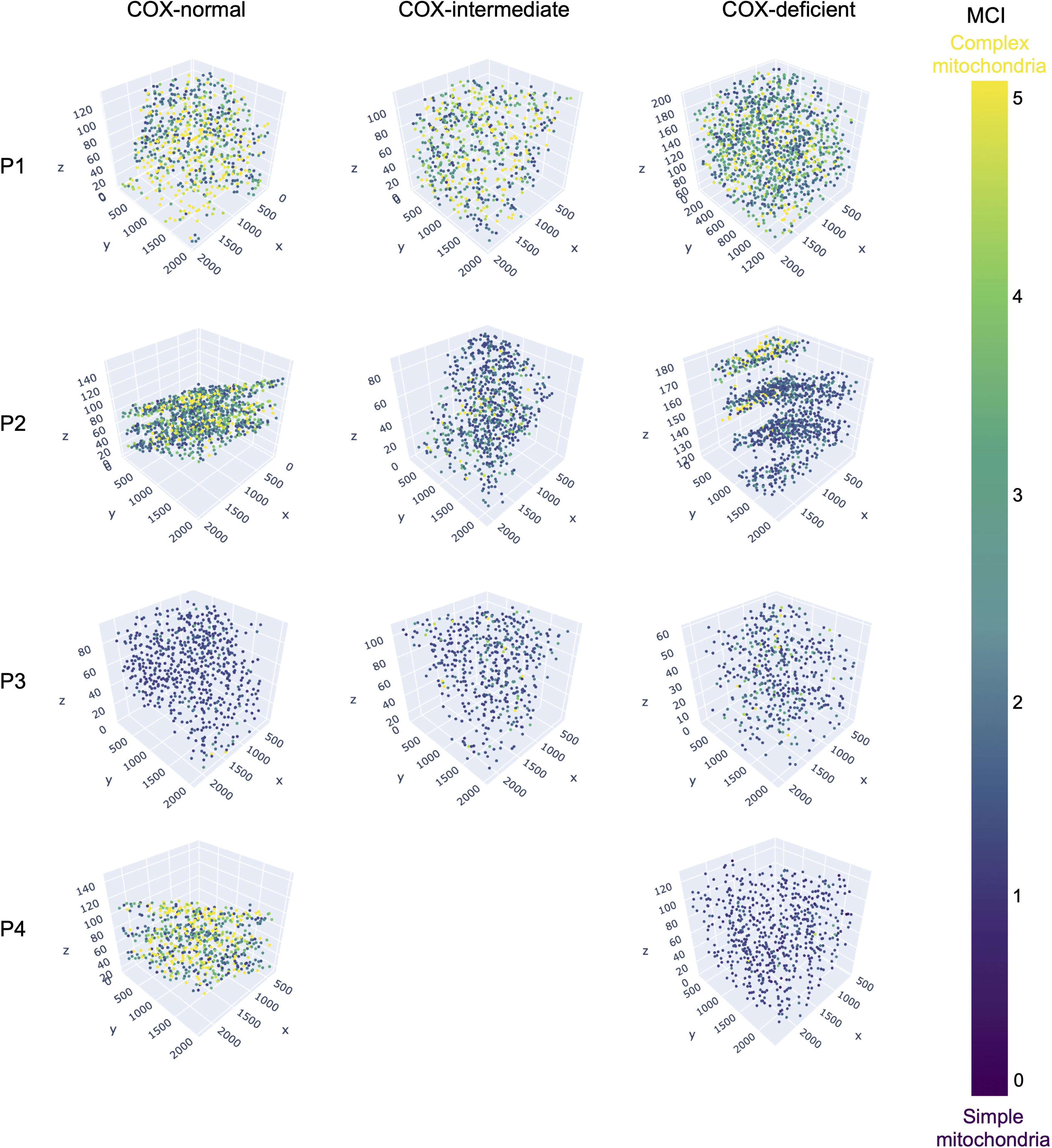
Mitochondrial spatial distribution and their MCI within a COX-normal/COX-intermediate and COX-deficient fibres from all patients. 3D scatter plot of mitochondrial complexity index across two sarcomeres in COX normal (A), COX-intermediate and COX-deficient fibres. Mitochondria are colour coded dependent on mitochondrial complexity index as indicated by the scale.

## Discussion

This study investigated the spatial distribution of COX activity and morphology of mitochondria in muscle biopsies of patients with mitochondrial myopathy throughout by volume-EM. Previous work in cultured cells has demonstrated that a relationship exists between mitochondrial morphology and function (Pich, Bach et al. 2005, Yu, Robotham and Yoon 2006, Benard, Bellance et al. 2007, Gomes, Di Benedetto and Scorrano 2011). In addition, in muscle biopsies from patients with mtDNA disease, mitochondria appeared damaged, more fragmented (Vincent, White et al. 2019) and also exhibited aberrant cristae (Vincent, Ng et al. 2016). However, it was necessary to have an assay capable of detecting both function and morphology, to be able to assess the relationship between these in patient tissue.

Using our COX-SBFSEM assay we present the analysis of both mitochondrial morphology and function in human muscle biopsies. These biopsies are from patients with single, large- scale mtDNA deletions, which will be present in individual muscle fibres at different levels of heteroplasmy, leading to a mosaic pattern of deficient and normal muscle fibres. Across all mitochondria presented here we generally find that lower COX intensities are associated with smaller mitochondrial complexity index (MCI) values and vice versa. Similar findings have been reported in our previous work where higher levels of m.8344A>G mutation in patient muscle are associated with more fragmented mitochondria, however due to the lack of COX activity measurements no direct correlation was possible in that study (Vincent, White et al. 2019). This also fits with previous reports from cybrids where a high level of mtDNA heteroplasmy which will be associated with mitochondrial dysfunction, leads to fragmentation of the mitochondrial network (Picard, Zhang et al. 2014). However, interestingly in the cybrids a medium level of mutation load which would be associated with lower levels of mitochondrial dysfunction are associated with more branched mitochondria. This is at odds with what we see here, however since all previous findings that have been able to observe mitochondrial function and morphology simultaneously have not used patient muscle samples, it is likely this is an important difference between the human muscle and cell models.

Another interesting observation from our ability to look in situ is that each fibre is in fact a mix of mitochondria with different COX activities. Confocal and electron microscopy have revealed heterogeneous COX activity and mitochondrial membrane potential, with an altered morphology for the COX deficient mitochondria in individual cells derived from clonal heteroplasmic cybrid cell lines with the m.3243A>G mutation (Bakker, Barthélémy et al. 2000). This fits with the mixture of COX activities we observe here in a single cell, since the mitochondrial membrane potential is related to the function of the respiratory chain and thus to mitochondrial morphology (Nicholls 2004, Picard, Shirihai et al. 2013). Vincent, White et al. (2019) also showed that mitochondrial morphological heterogeneity is demonstrated by the coexistence of a wide variation in morphology within a cell or patient (Vincent, White et al. 2019). The presence of a moderate level of mtDNA heteroplasmy in mitochondrial disease was associated with more heterogeneity (Vincent, White et al. 2019). Here, the results indicate a heterogeneous population of complex and simple mitochondria inter- or intracellularly in patient muscles. Interestingly the same pattern can be seen in all patients even if they harbour different deletions.

Within the COX-deficient fibres, mitochondria appeared to be simpler with a lower MCI in all patients, whereas COX-normal fibres exhibit a higher MCI. However, the pattern observed in COX deficient, intermediate and normal fibres differs. The mitochondrial fragmentation exhibited in COX-deficient fibres, suggest that the dysfunction is too high and there’s may no longer sufficient energy available to carry out fusion. Alternatively, in fibres with low levels of dysfunction, such as COX-normal and intermediate fibres, mitochondrial fusion may restore homeostasis, promote mtDNA mixing and act as a buffer for mitochondrial dysfunction (Chen, Vermulst et al. 2010).

Within a segment of skeletal muscle fibre, a proportion of mutated mtDNA may exceed a critical threshold, resulting in COX deficiency (Seligman, Karnovsky et al. 1968). When mtDNA variants reach a high mutation load due to intracellular clonal expansion, a biochemical defect results in COX deficient segments, contributing to the appearance of clinical pathology and disease progression (Schon, Bonilla and DiMauro 1997). Chinnery, Howel et al. (2003) provide objective evidence that the clinical progression of mtDNA myopathy is associated with a biochemical deficiency that develops independently within individual muscle fibres. To understand how the mtDNA variant proliferate and the spread of the deficiency during clonal expansion, Vincent, Rosa et al. (2018) studied muscle fibres from patients with mitochondrial disease. The results from this study showed deficiency to occur in focal regions surrounding the myonuclei, inducing mitochondrial biogenesis to proliferate before spreading through the muscle fibre. The deficient mitochondria will propagate transversely first via direct physical interactions between mitochondria before propagating longitudinally where fewer mitochondrial connections are observed (Vincent, Rosa et al. 2018, Vincent, White et al. 2019).

The EM data obtained here matches with previous reports of segmental COX deficiency within cells (Larsson and Oldfors 2001, Vincent, Rosa et al. 2018). What is particularly interesting is that in normal fibres small numbers of isolated COX deficient mitochondria are observed. In intermediate fibres in comparison, we see a mix of mitochondrial activities and morphologies and no real spatial pattern. This suggests to us that it is this intermediate stage where the spread of dysfunction is most likely to occur (Figure 10). This would fit with previous findings (Picard, Zhang et al. 2014, Vincent, White et al. 2019) which demonstrate that at low levels of mtDNA heteroplasmy mitochondria are more branched and connected and therefore there is a greater opportunity to share material and for dysfunction to spread. However, what is interesting to us in the COX normal muscle fibres that we have examined is that there appear to be small numbers of negative mitochondria with a strong spatial segregation of COX activity and a small region of intermediate mitochondria that act as a transition zone between the normal and deficient mitochondria (Figure 10). These fibres we believe to be important to informing our knowledge of how dysfunction spreads, with the intermediate mitochondria being the key players. The COX deficient mitochondria have become isolated from the network but presumably too late as they have likely already shared some degree of dysfunction with the adjacent mitochondria which are now intermediate mitochondria branching and interacting with the COX-normal mitochondria.

**Figure 10.**
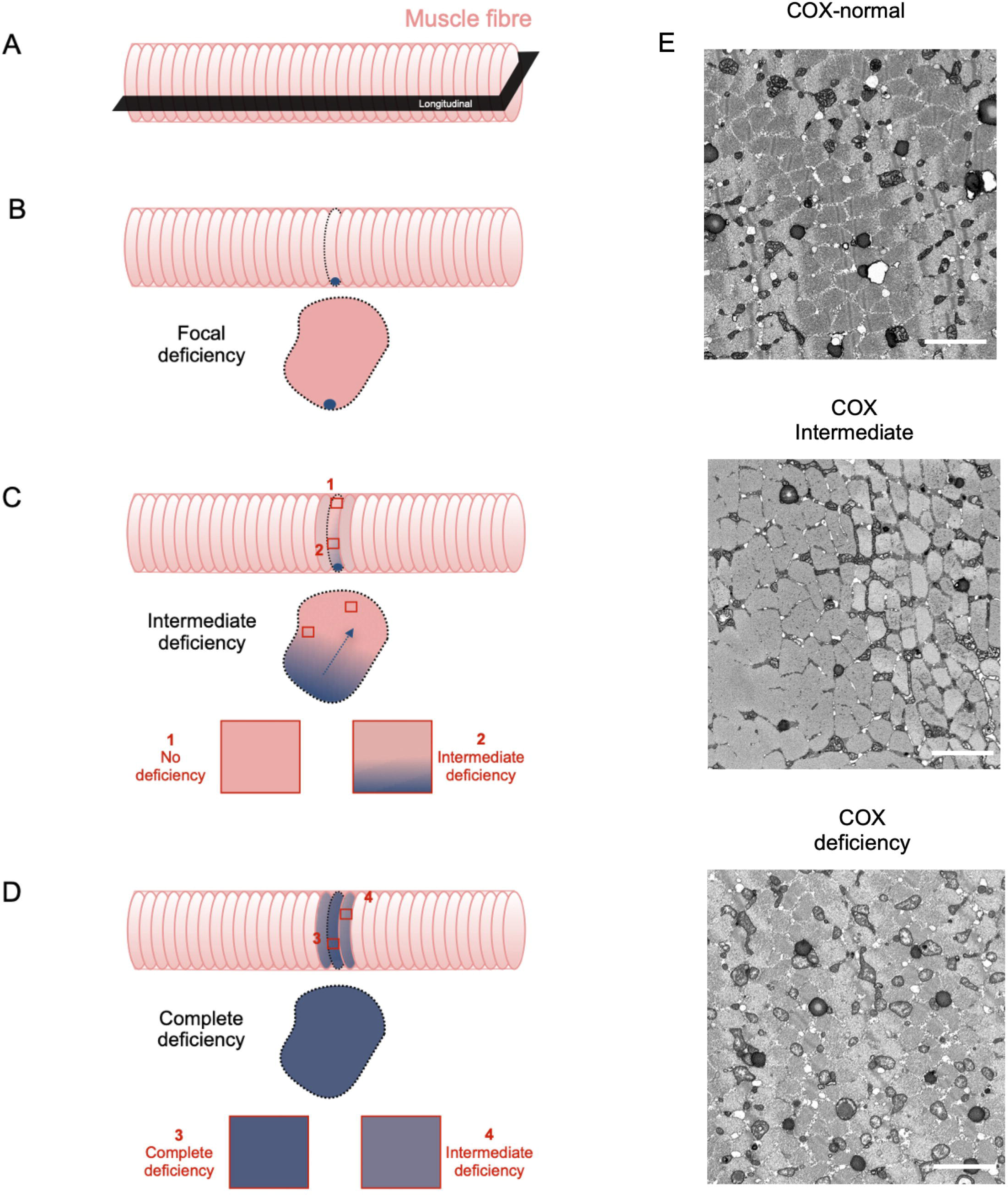
Model illustrating spread of COX-deficiency. **(A-F)** A theoretical guide to understanding and classifying the development of COX deficiency. Focal perinuclear deficiency **(B)** to segmental deficiency **(C).** The spread of the dysfunction involves COX-intermediate stages before the full complete COX-deficiency muscle fibre **(D).** Examples of 2D images from Fibres COX-normal, intermediate and deficient **(E).**

The findings of Murphy, Ratnaike et al. (2012), using COX histochemistry, have demonstrated that intermediate levels of COX activity exist between COX-normal regions and fully COX-deficient segments of muscle fibres, which they term as COX-intermediate segments. This COX-intermediate segment effectively being the transition zone, between normal and deficient regions, where dysfunction is spreading along the fibre. Thus, this diverse range of mitochondrial COX activity could underlie how the COX-deficiency within individual muscle fibres ranges from focal to segmental and complete deficiency in patients with mtDNA disease (Johnson, Turnbull et al. 1983, Vincent, Rosa et al. 2018)(Figure 10).

In this work we have aimed to understand the link between mitochondrial function and mitochondrial morphology *in situ*, however it is worth noting an important limitation. Patients with mtDNA deletions will have deletions that remove different numbers of genes encoding subunits for complexes I, III, IV and V as well as tRNAs. As such it is possible that whilst our assay shows a muscle fibre or even a mitochondrion as being COX positive it may have a deficiency in one of the other complexes which we are unable to detect with this assay. However, the fact that we see clear differences between the different muscle fibre classes suggests to us this is a have only a small impact on the fibres analysed here.

A limitation here is that each ROI was imaged from the centre of a fibre. Therefore, a strong spatial pattern cannot easily be observed, as it is just a part of each individual fibre that is imaged and only a small depth through the fibre. However, these results may suggest that each fibre is at a particular stage of COX deficiency, due to the segmental spread of the dysfunction. Furthermore, for some fibres, the clusters were well defined, especially with COX-normal, and intermediate fibres along the muscle fibre diameter. This is important in terms of progression and assessing mitochondrial disease severity. Alongside this it is challenging to determine the type of muscle fibre type using electron microscopy, as the quadriceps muscle contains a mixture of muscle fibres with different properties. This can be problematic because muscle fibres exhibit different mitochondrial volume density and morphology. In the future, detecting fibre types would allow for a better understanding. However, the number of investigated fibres from each patient is a compromise between exploring enough to account for variability and feasibility due to the manual work required for accurate mitochondrial segmentation.

## Conclusions

This work provided the first quantitative assessment in single, large-scale mtDNA deletion patients’ skeletal muscle of mitochondrial morphology and COX function using SBFSEM. This study shows first the spectrum of COX activity within muscle fibres, with three distinct mitochondrial populations: normal, intermediate, and deficient. Secondly, it shows the distribution of mitochondrial dysfunction and disease progression through COX deficient and normal fibres. It is also interesting to observe some degree of spatial pattern, and this warrants further investigation. Finally, this study gives new insights into how mitochondrial function is linked to mitochondrial morphology in situ. The mitochondria in COX- intermediate fibres exhibit activity levels between those seen in COX-normal fibres and those that are COX-deficient. The COX-normal fibres exhibited complex mitochondria compared to spherical mitochondria within COX-deficient fibres.

## Acknowledgements

The work was supported by the Wellcome Centre for Mitochondrial Research, NIHR Biomedical Research Centre and used a Gatan 3view purchased through a BBSRC grant (BB/M012093/1). AEV was supported by a Sir Henry Wellcome Postdoctoral Fellowship (215888/Z/19/Z) and Newcastle University Academic Track Fellowship. JF was supported by a Wellcome Ph.D. project match funded by Newcastle University awarded to (C0163N3031). We would like to thank the AIMM Trial Participants and AIMM Trial Group for providing and collecting the muscle samples for this work.

## Author Contributions

JF, AEV and DMT designed the study. JF, TD and RL developed the methodology. JF and TD prepared the samples. JF, TD and RL imaged the samples. JF analysed EM images. HT and EF collected genetic data for the samples. JF and CL performed data analysis. JF, CL and AEV interpreted the data. JF and AEV wrote the manuscript draft. All authors reviewed and approved the manuscript.

## Data Availability

The raw data image data, models and a spreadsheet of quantitative data have been deposited here and available to download [link].

## References

1. Bakker, A., C. Barthélémy, P. Frachon, D. Chateau, D. Sternberg, J. P. Mazat and A. Lombès (2000). “Functional Mitochondrial Heterogeneity in Heteroplasmic Cells Carrying the Mitochondrial DNA Mutation Associated with the MELAS Syndrome (Mitochondrial Encephalopathy, Lactic Acidosis, and Strokelike Episodes).” Pediatric Research 48(2): 143–150.

2. Belevich, I., M. Joensuu, D. Kumar, H. Vihinen and E. Jokitalo (2016). “Microscopy Image Browser: A Platform for Segmentation and Analysis of Multidimensional Datasets.” PLoS Biol 14(1): e1002340.

3. Benard, G., N. Bellance, D. James, P. Parrone, H. Fernandez, T. Letellier and R. Rossignol (2007). “Mitochondrial bioenergetics and structural network organization.” J Cell Sci 120(Pt 5): 838–848.

4. Chen, H., M. Vermulst, Y. E. Wang, A. Chomyn, T. A. Prolla, J. M. McCaffery and D. C. Chan (2010). “Mitochondrial fusion is required for mtDNA stability in skeletal muscle and tolerance of mtDNA mutations.” Cell 141(2): 280–289.

5. Chinnery, P. F., D. Howel, D. M. Turnbull and M. A. Johnson (2003). “Clinical progression of mitochondrial myopathy is associated with the random accumulation of cytochrome c oxidase negative skeletal muscle fibres.” J Neurol Sci 211(1-2): 63–66.

6. Dimauro, S., J. F. Nicholson, A. P. Hays, A. B. B, A. Papadimitriou, R. Koenigsberger and D. C. Devivo (1983). “Benign infantile mitochondrial myopathy due to reversible cytochrome c oxidase deficiency.” Annals of Neurology 14(2): 226–234.

7. DiMauro, S., M. Zeviani, E. Bonilla, N. Bresolin, M. Nakagawa, A. F. Miranda and M. Moggio (1985). “Cytochrome c oxidase deficiency.” Biochem Soc Trans 13(4): 651–653.

8. Eisner, D. A., J. L. Caldwell, K. Kistamás and A. W. Trafford (2017). “Calcium and Excitation-Contraction Coupling in the Heart.” Circ Res 121(2): 181–195.

9. Faitg, J., T. Davey, D. M. Turnbull, K. White and A. E. Vincent (2020). “Mitochondrial morphology and function: two for the price of one!” Journal of Microscopy 278(2): 89–106.

10. Gomes, L. C., G. Di Benedetto and L. Scorrano (2011). “During autophagy mitochondria elongate, are spared from degradation and sustain cell viability.” Nat Cell Biol 13(5): 589–598.

11. Johnson, M., D. Turnbull, D. Dick and H. Sherratt (1983). “A partial deficiency of cytochrome c oxidase in chronic progressive external ophthalmoplegia.” Journal of the neurological sciences 60(1): 31–53.

12. Larsson, N. G. and A. Oldfors (2001). “Mitochondrial myopathies.” Acta Physiol Scand 171(3): 385–393.

13. Morgan-Hughes, J. A. (1986). “Mitochondrial diseases.” Trends in Neurosciences 9: 15–19.

14. Murphy, J. L., T. E. Ratnaike, E. Shang, G. Falkous, E. L. Blakely, C. L. Alston, T. Taivassalo, R. G. Haller, R. W. Taylor and D. M. Turnbull (2012). “Cytochrome c oxidase-intermediate fibres: importance in understanding the pathogenesis and treatment of mitochondrial myopathy.” Neuromuscul Disord 22(8): 690–698.

15. Nicholls, D. G. (2004). “Mitochondrial membrane potential and aging.” Aging Cell 3(1): 35–40.

16. Nonaka, I., Y. Koga, E. Ohtaki and M. Yamamoto (1989). “Tissue specificity in cytochrome c oxidase deficient myopathy.” J Neurol Sci 92(2-3): 193–203.

17. Ogata, T. and Y. Yamasaki (1985). “Scanning electron-microscopic studies on the three- dimensional structure of mitochondria in the mammalian red, white and intermediate muscle fibers.” Cell and Tissue Research 241(2): 251–256.

18. Picard, M., R. T. Hepple and Y. Burelle (2012). “Mitochondrial functional specialization in glycolytic and oxidative muscle fibers: tailoring the organelle for optimal function.” Am J Physiol Cell Physiol 302(4): C629–641.

19. Picard, M., O. S. Shirihai, B. J. Gentil and Y. Burelle (2013). “Mitochondrial morphology transitions and functions: implications for retrograde signaling?” American Journal of Physiology-Regulatory, Integrative and Comparative Physiology 304(6): R393–R406.

20. Picard, M., K. White and D. M. Turnbull (2013). “Mitochondrial morphology, topology, and membrane interactions in skeletal muscle: a quantitative three-dimensional electron microscopy study.” J Appl Physiol (1985) 114(2): 161–171.

21. Picard, M., J. Zhang, S. Hancock, O. Derbeneva, R. Golhar, P. Golik, S. O’Hearn, S. Levy, P. Potluri, M. Lvova, A. Davila, C. S. Lin, J. C. Perin, E. F. Rappaport, H. Hakonarson, I. A. Trounce, V. Procaccio and D. C. Wallace (2014). “Progressive increase in mtDNA 3243A>G heteroplasmy causes abrupt transcriptional reprogramming.” Proceedings of the National Academy of Sciences 111(38): E4033–E4042.

22. Pich, S., D. Bach, P. Briones, M. Liesa, M. Camps, X. Testar, M. Palacín and A. Zorzano (2005). “The Charcot-Marie-Tooth type 2A gene product, Mfn2, up-regulates fuel oxidation through expression of OXPHOS system.” Hum Mol Genet 14(11): 1405–1415.

23. Ranieri, M., S. Brajkovic, G. Riboldi, D. Ronchi, F. Rizzo, N. Bresolin, S. Corti and G. P. Comi (2013). “Mitochondrial fusion proteins and human diseases.” Neurol Res Int 2013: 293893.

24. Rocha, M. C., H. S. Rosa, J. P. Grady, E. L. Blakely, L. He, N. Romain, R. G. Haller, J. Newman, R. McFarland, Y. S. Ng, G. S. Gorman, A. M. Schaefer, H. A. Tuppen, R. W. Taylor and D. M. Turnbull (2018). “Pathological mechanisms underlying single large-scale mitochondrial DNA deletions.” Ann Neurol 83(1): 115–130.

25. Rossignol, R., B. Faustin, C. Rocher, M. Malgat, J. P. Mazat and T. Letellier (2003). “Mitochondrial threshold effects.” Biochem J 370(Pt 3): 751–762.

26. Schon, E. A., E. Bonilla and S. DiMauro (1997). “Mitochondrial DNA Mutations and Pathogenesis.” Journal of Bioenergetics and Biomembranes 29(2): 131–149.

27. Seligman, A. M., M. J. Karnovsky, H. L. Wasserkrug and J. S. Hanker (1968). “Nondroplet ultrastructural demonstration of cytochrome oxidase activity with a polymerizing osmiophilic reagent, diaminobenzidine (DAB).” J Cell Biol 38(1): 1–14.

28. Vincent, A. E., Y. S. Ng, K. White, T. Davey, C. Mannella, G. Falkous, C. Feeney, A. M. Schaefer, R. McFarland, G. S. Gorman, R. W. Taylor, D. M. Turnbull and M. Picard (2016). “The Spectrum of Mitochondrial Ultrastructural Defects in Mitochondrial Myopathy.” Sci Rep 6: 30610.

29. Vincent, A. E., H. S. Rosa, K. Pabis, C. Lawless, C. Chen, A. Grünewald, K. A. Rygiel, M. C. Rocha, A. K. Reeve, G. Falkous, V. Perissi, K. White, T. Davey, B. J. Petrof, A. A. Sayer, C. Cooper, D. Deehan, R. W. Taylor, D. M. Turnbull and M. Picard (2018). “Subcellular origin of mitochondrial DNA deletions in human skeletal muscle.” Ann Neurol 84(2): 289–301.

30. Vincent, A. E., K. White, T. Davey, J. Philips, R. T. Ogden, C. Lawless, C. Warren, M. G. Hall, Y. S. Ng, G. Falkous, T. Holden, D. Deehan, R. W. Taylor, D. M. Turnbull and M. Picard (2019). “Quantitative 3D Mapping of the Human Skeletal Muscle Mitochondrial Network.” Cell Rep 27(1): 321.

31. Wilke, S. A., J. K. Antonios, E. A. Bushong, A. Badkoobehi, E. Malek, M. Hwang, M. Terada, M. H. Ellisman and A. Ghosh (2013). “Deconstructing complexity: serial block-face electron microscopic analysis of the hippocampal mossy fiber synapse.” J Neurosci 33(2): 507–522.

32. Yu, T., J. L. Robotham and Y. Yoon (2006). “Increased production of reactive oxygen species in hyperglycemic conditions requires dynamic change of mitochondrial morphology.” Proc Natl Acad Sci U S A 103(8): 2653–2658.

