## Supplementary Figures 1-7 for "Mitochondrial morphology and function in mitochondrial disease"

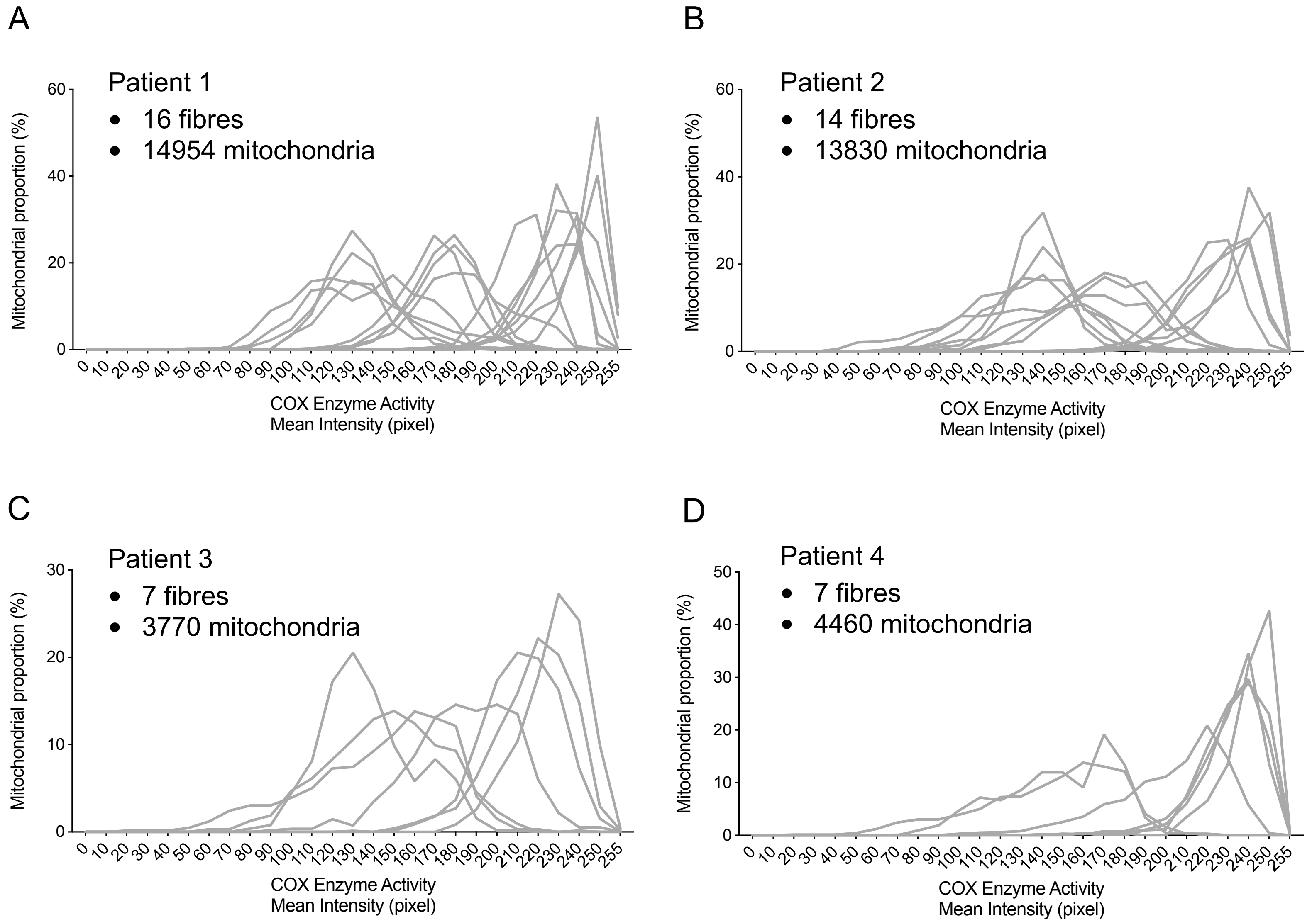


**Figure S1. Mitochondrial COX activity for individual fibres.**

Frequency distribution of mitochondrial COX activity from individual fibres of single, large-scale mtDNA deletion Patient 1 **(A),** Patient 2 **(B),** Patient 3 **(C)** and Patient 4 **(D).** The selection of the fibres was assessed by eyes. For patient 1, 16 fibres were analysed with a total of 14954 mitochondria and for Patient 2, 13 fibres were analysed with a total of 13830 mitochondria, for Patient 3, 7 fibres were analysed with a total of 3770 mitochondria and for Patient 4, 7 fibres were analysed with a total of 4460 mitochondria analysed.

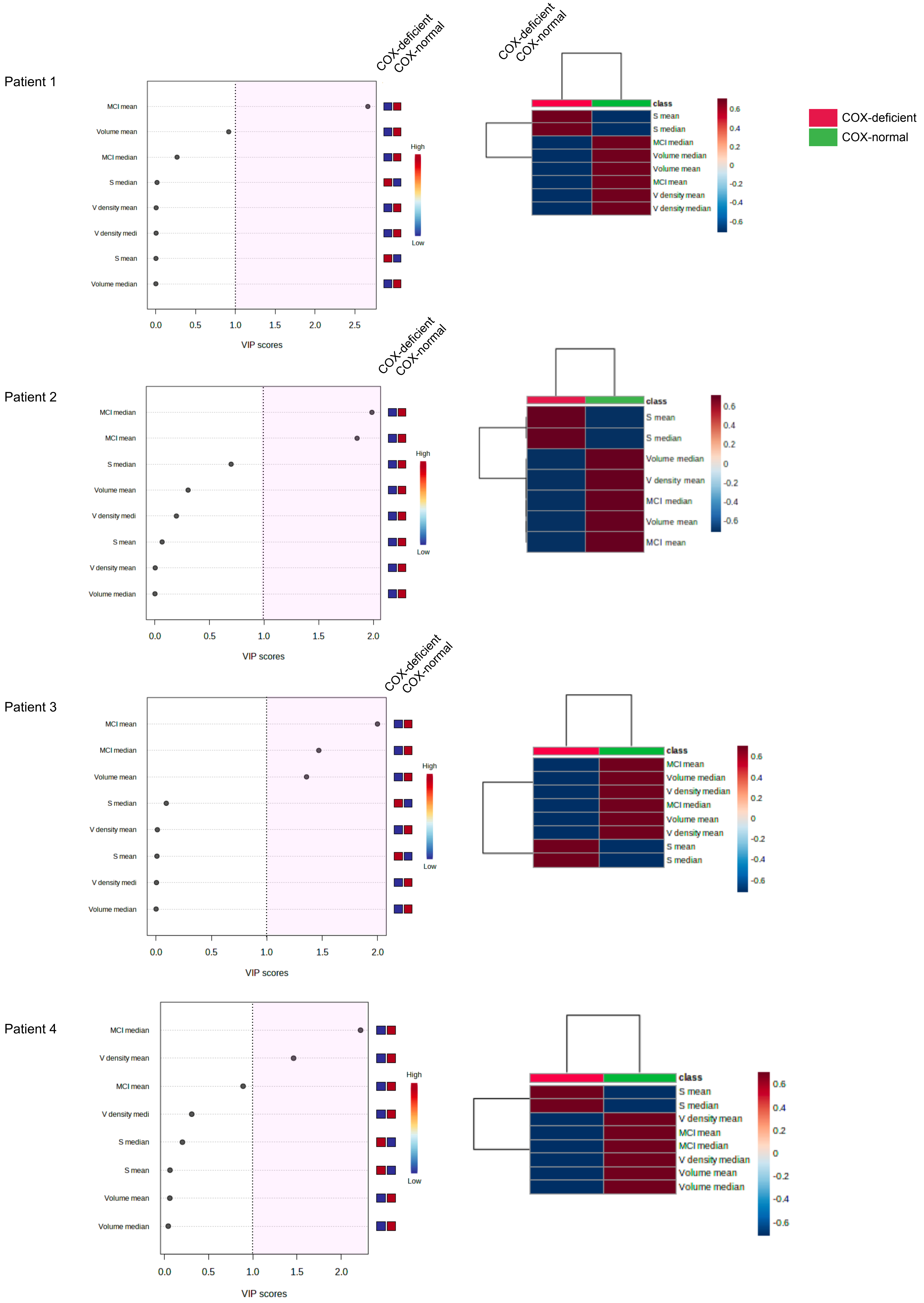


**Figure S2. Multivariate analysis of mitochondrial morphology between individual normal and deficient fibres from patient 1,2,3 & 4.**

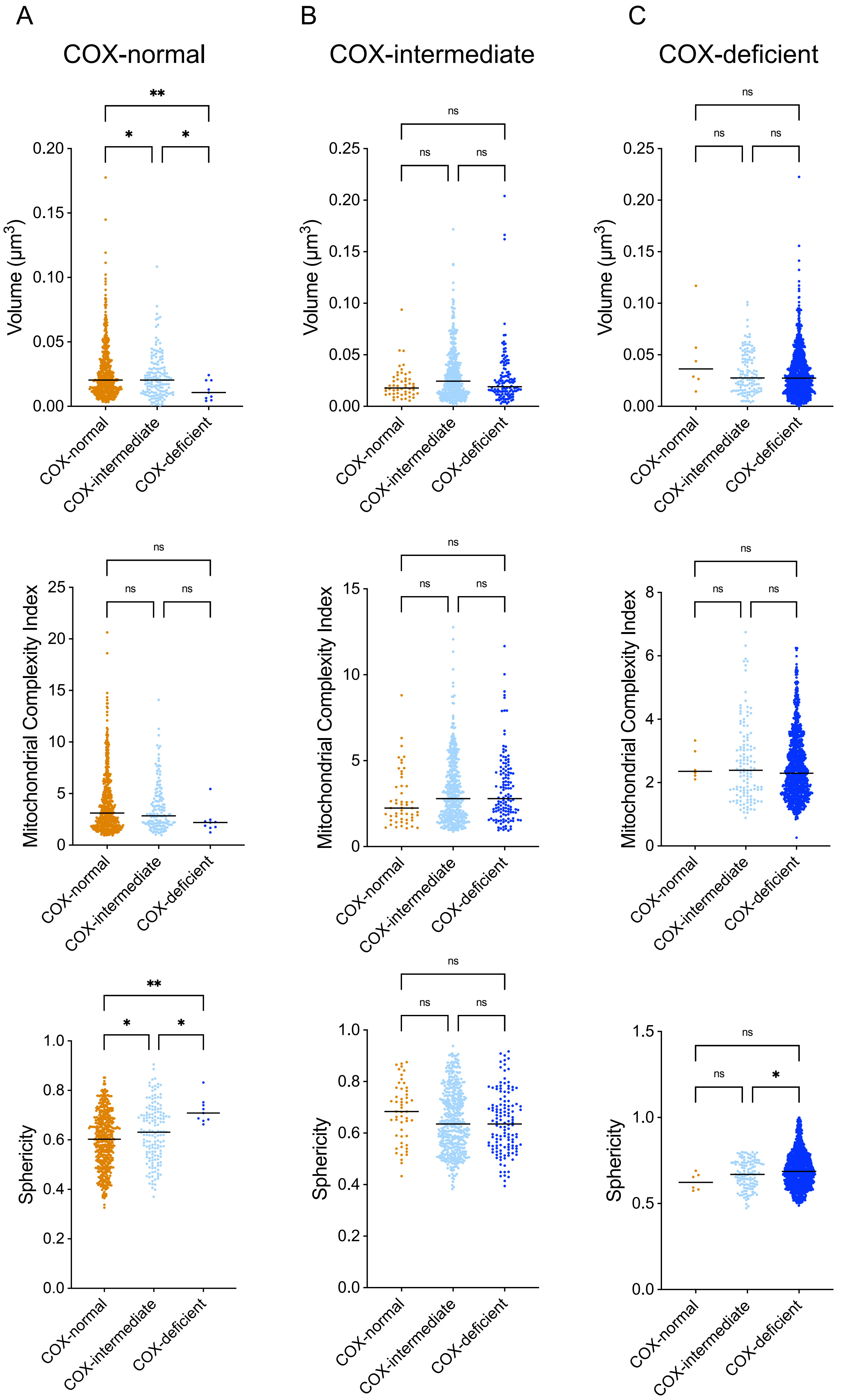


**Figure S3. Mitochondrial morphology comparison from COX-normal, intermediate, and deficient fibres from Patient 1.**
Scatter plot showing mitochondrial volume **(A)**, MCI **(B)** and sphericity **(C)** of COX-normal, COX-intermediate and COX-deficient mitochondria from respective fiber

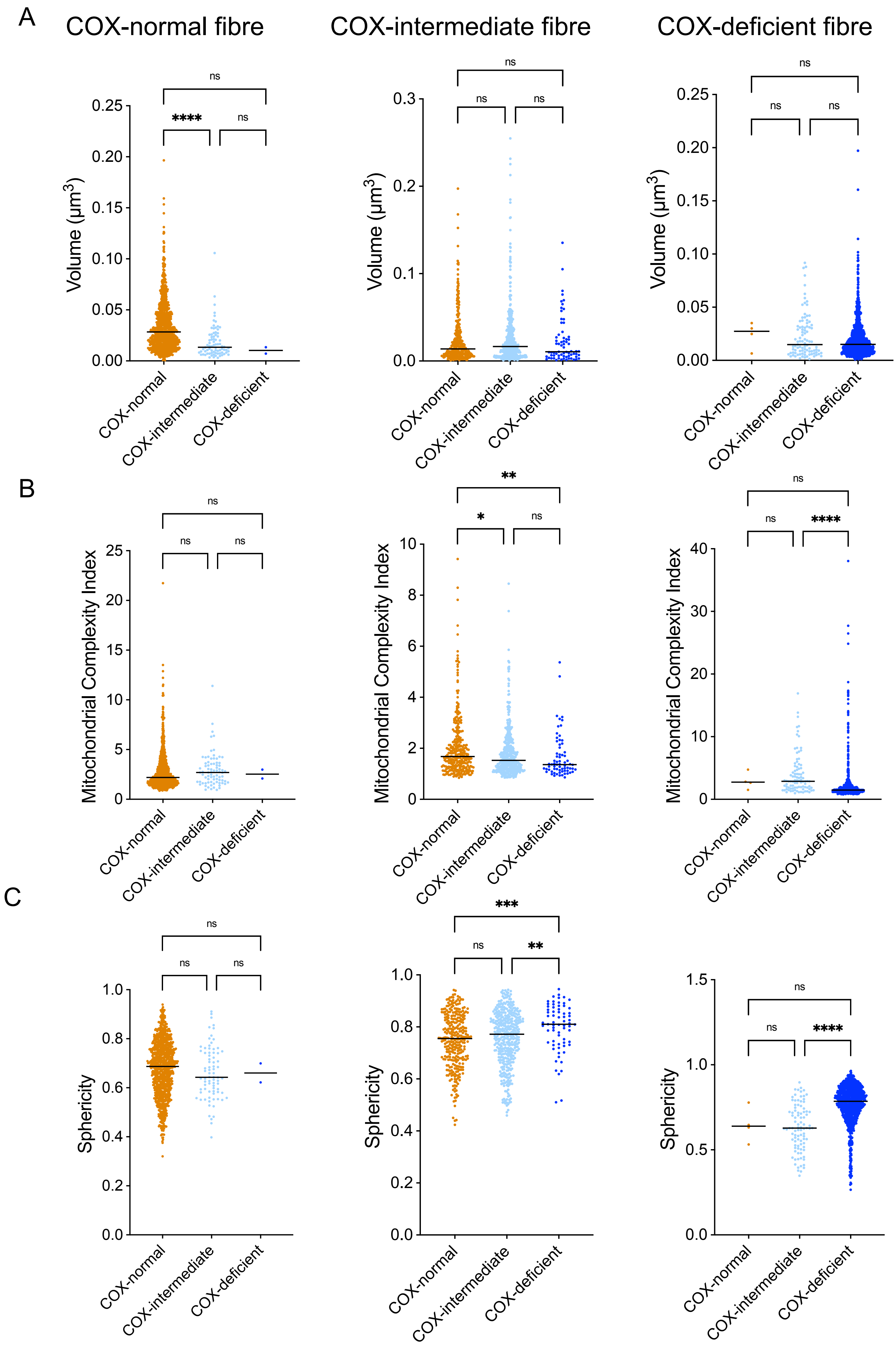


**Figure S4. Mitochondrial morphology comparison from COX-normal, intermediate, and deficient fibres from Patient 2**
Scatter plot showing mitochondrial volume **(A)**, MCI **(B)** and sphericity **(C)** of COX-normal, COX-intermediate and COX-deficient mitochondria from respective fiber

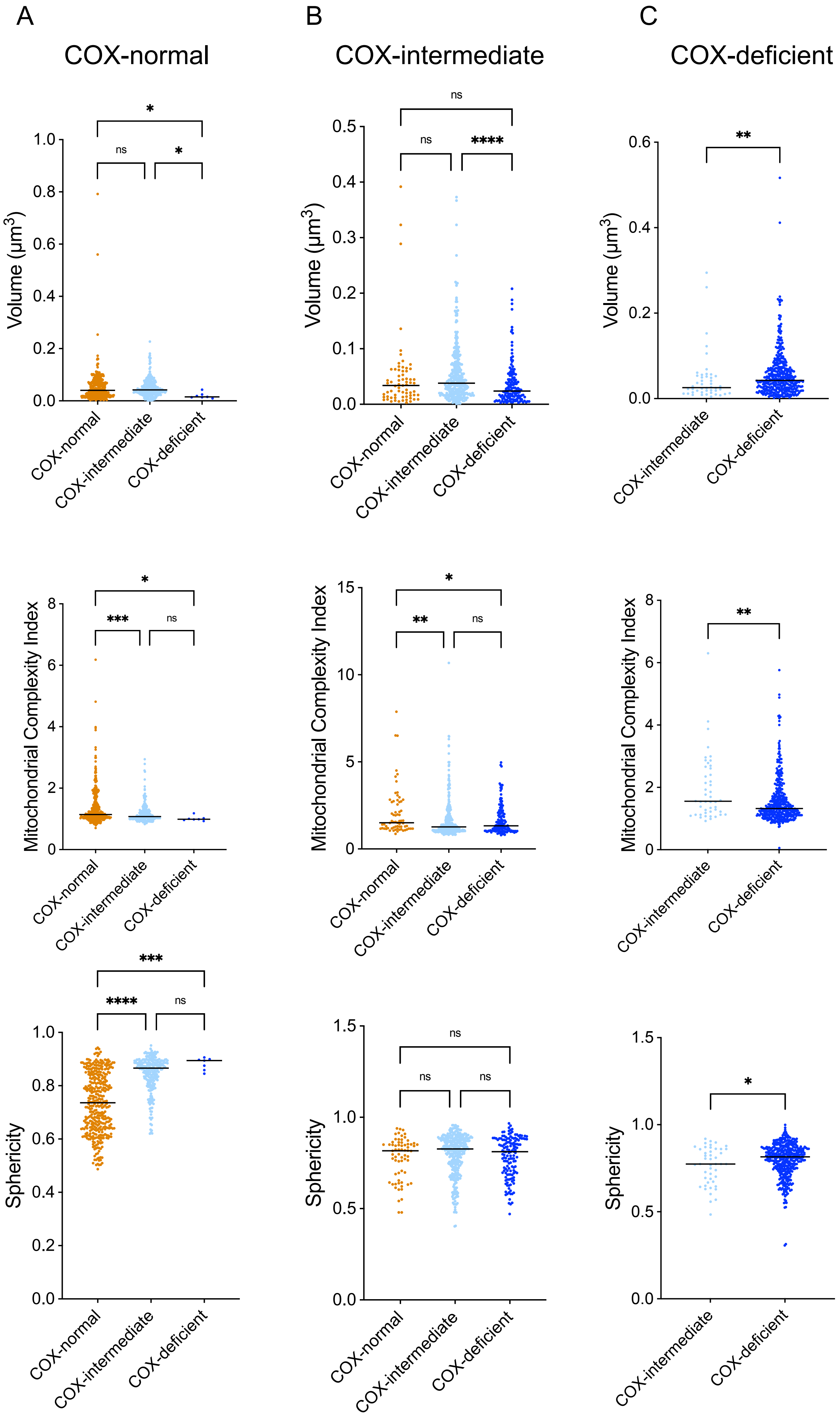


**Figure S5. Mitochondrial morphology comparison from COX-normal, intermediate, and deficient fibres from Patient 3**
Scatter plot showing mitochondrial volume **(A)**, MCI **(B)** and sphericity **(C)** of COX-normal, COX-intermediate and COX-deficient mitochondria from respective fiber

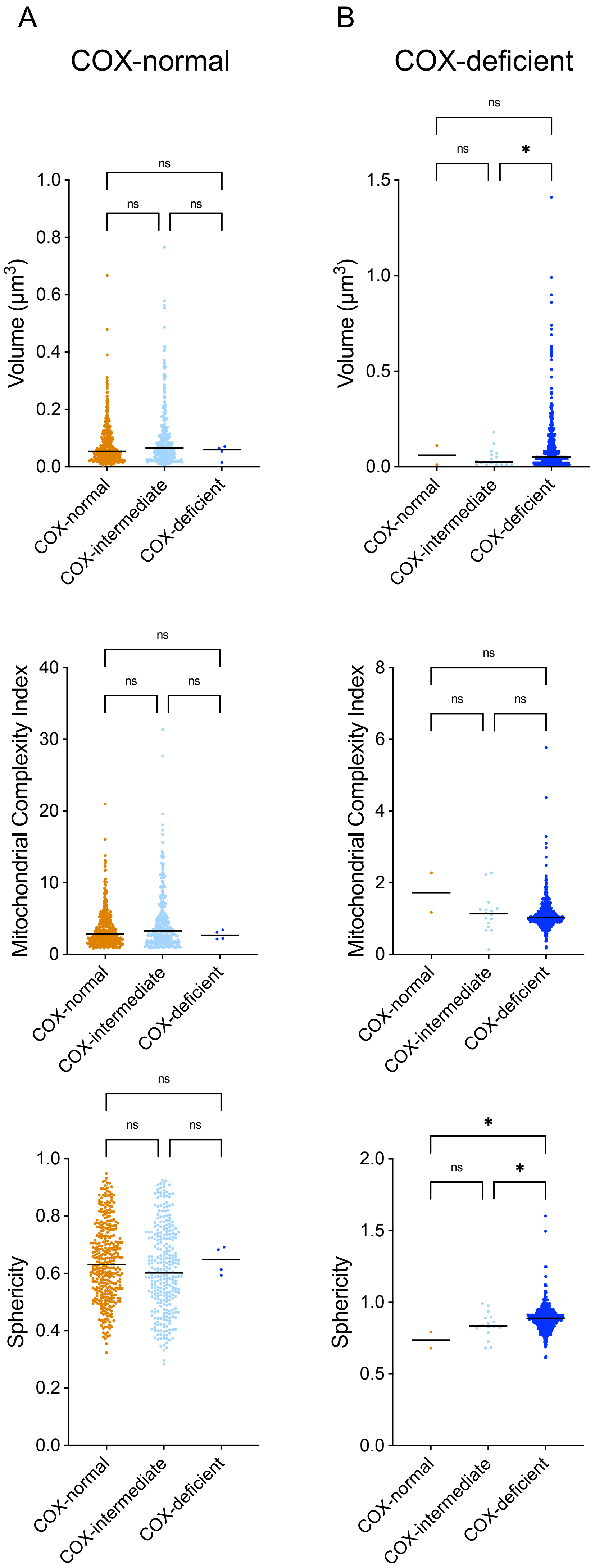


**Figure S6. Mitochondrial morphology comparison from COX-normal, intermediate, and deficient fibres from Patient 4**
Scatter plot showing mitochondrial volume **(A)**, MCI **(B)** and sphericity **(C)** of COX-normal, COX-intermediate and COX-deficient mitochondria from respective fiber

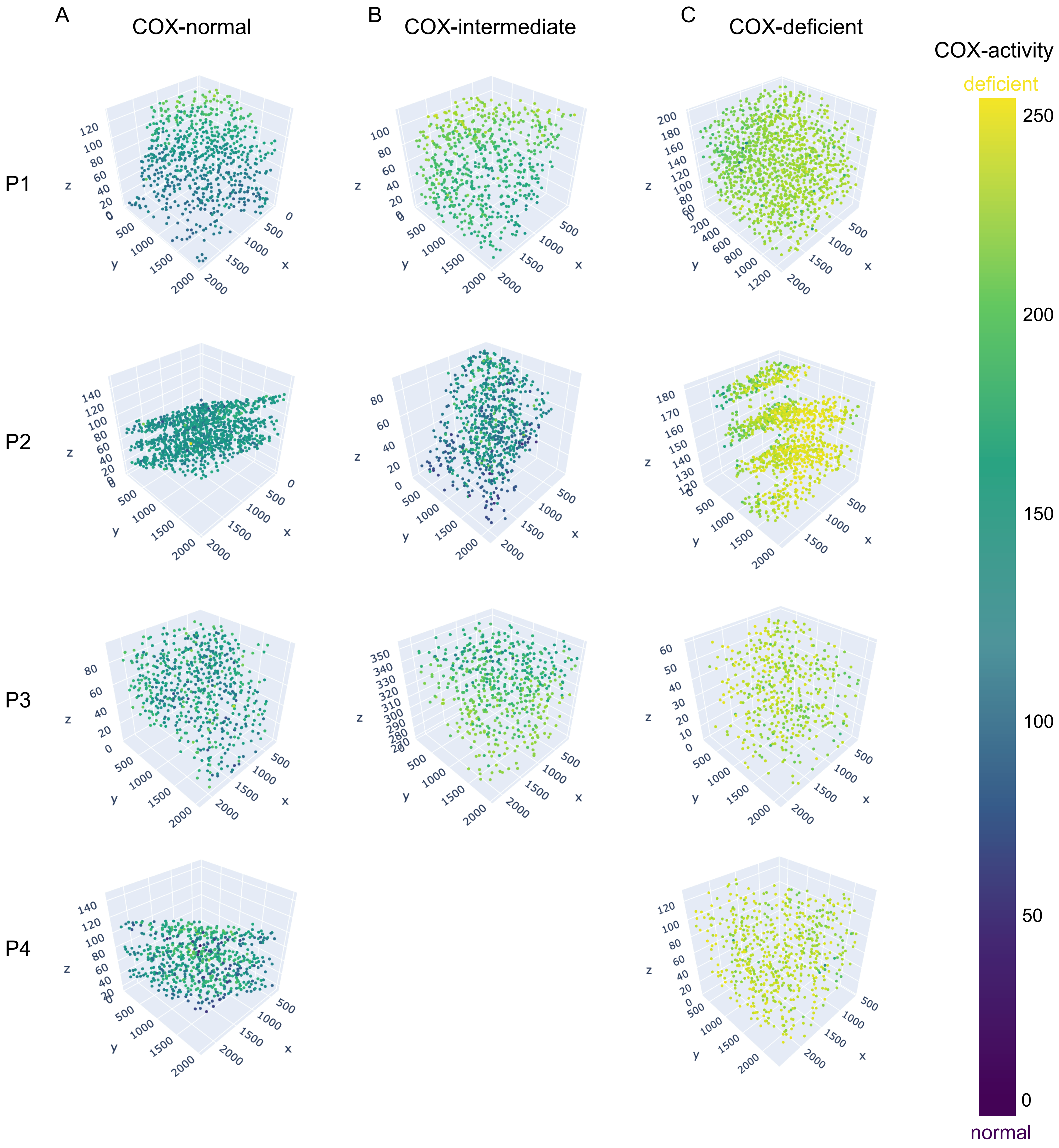


**Figure S7. COX activity and spatial distribution of the mitochondrial COX activity and their respective morphologies within a COX-normal/COX-intermediate and COX-deficient fibres from Patient 1,2,3&4 from figure 8**

The 3D scatter plot of a COX-normal (Figure 7A), COX-intermediate (Figure 7B) and COX-deficient (Figure 7C) fibre demonstrate coloured mitochondria according to their COX activity.

3D scatter plot of mitochondrial COX activity across two sarcomeres in COX normal (A), COX-intermediate (B) and COX-deficient (C) fibres demonstrate coloured mitochondria according to their COX activity.

Mitochondria are colour coded dependent on COX activity as indicated by the scale Cluster of COX-normal and intermediate mitochondria can be distinguished by the dotted circle.
