## Supplementary Tables for "Mitochondrial morphology and function in mitochondrial disease"

| Patients | Fibre ID | COX normal (%) | COX intermediate (%) | COX deficient (%) |
| --- | --- | --- | --- | --- |
| Patient1 | Fibre 1 | 82.3 | 17.4 | 0.3 |
|  | Fibre 8 | 71.6 | 27.9 | 0.4 |
|  | Fibre 13 | 92.5 | 7.5 | 0 |
|  | Fibre 16 | 62.3 | 37.7 | 0 |
|  | Fibre 14 | 87.8 | 12.2 | 0 |
|  | Fibre 2 | 8.8 | 88.8 | 2.4 |
|  | Fibre 9 | 7.5 | 91.9 | 0.6 |
|  | Fibre 10 | 5.5 | 79.6 | 14.9 |
|  | Fibre 15 | 13.9 | 85 | 1.1 |
|  | Fibre 3 | 0.5 | 22.1 | 77.4 |
|  | Fibre 4 | 0.1 | 10.5 | 89.4 |
|  | Fibre 5 | 0.1 | 6.6 | 93.3 |
|  | Fibre 6 | 0 | 0.8 | 99.2 |
|  | Fibre 7 | 0 | 14.2 | 85.8 |
|  | Fibre 11 | 0 | 41.7 | 58.3 |
|  | Fibre 12 | 0 | 41.7 | 58.3 |
| Patient 2 | Fibre 1 | 54.5 | 44.6 | 0.8 |
|  | Fibre 5 | 69.4 | 29.4 | 1.2 |
|  | Fibre 6 | 85.6 | 14.3 | 0.1 |
|  | Fibre 7 | 74.2 | 25.8 | 0 |
|  | Fibre 12 | 84 | 15.9 | 0.1 |
|  | Fibre 2 | 15.8 | 80.2 | 4 |
|  | Fibre 4 | 24.3 | 74.3 | 1.5 |
|  | Fibre 11 | 34.5 | 61.7 | 3.9 |
|  | Fibre 3 | 0.1 | 18.2 | 81.7 |
|  | Fibre 8 | 0.5 | 29.8 | 69.7 |
|  | Fibre 9 | 0.2 | 18.2 | 81.6 |
|  | Fibre 10 | 0.4 | 11.6 | 88 |
|  | Fibre 13 | 0.2 | 5.2 | 94.7 |
| Fibre 3 | Fibre 4 | 60.32 | 36.22 | 3.46 |
|  | Fibre 6 | 54.78 | 43.1 | 2.12 |
|  | Fibre 7 | 77.8 | 21.8 | 0.4 |
|  | Fibre 3 | 12.23 | 60.95 | 26.82 |
|  | Fibre 1 | 0 | 9.95 | 90.05 |
|  | Fibre 2 | 0.42 | 23.22 | 76.36 |
|  | Fibre 5 | 0.26 | 16.82 | 82.92 |
| Fibre 4 | Fibre 6 | 56.80 | 45.2 | 2 |
|  | Fibre 7 | 75.2 | 23.3 | 1.4 |
|  | Fibre 1 | 6.59 | 26.44 | 66.97 |
|  | Fibre 2 | 0.25 | 2.4 | 97.35 |
|  | Fibre 5 | 0 | 0.8 | 99.2 |
|  | Fibre 4 | 0.33 | 1.65 | 98.02 |
|  | Fibre 3 | 0.44 | 3.65 | 95.91 |

Table S1 Percentage class of individual mitochondria in fibre of patients P1,P2,P3 & P4.

|  | **P1** | | |  | **P2** | | |  | **P3** | | |  | **P4** | |
| --- | --- | --- | --- | --- | --- | --- | --- | --- | --- | --- | --- | --- | --- | --- |
|  | F+ | F± | F- |  | F+ | F± | F- |  | F+ | F± | F- |  | F+ | F- |
| V < 10% | 10,15 | 20,58 | 30,91 |  | 10,05 | 14,19 | 26,44 |  | 10,05 | 15,15 | 15,47 |  | 9,91 | 11,17 |
| 10% < V < 90% | 79,87 | 78,51 | 68,54 |  | 79,98 | 71,75 | 69,43 |  | 79,97 | 84,85 | 84,12 |  | 80,18 | 83,29 |
| V > 90% | 9,97 | 0,91 | 0,55 |  | 9,97 | 14,06 | 4,13 |  | 9,98 | 0,00 | 0,41 |  | 9,91 | 5,54 |
| MCI < 10% | 10,03 | 30,86 | 43,51 |  | 10,01 | 13,74 | 31,29 |  | 9,98 | 5,66 | 14,49 |  | 9,91 | 66,39 |
| 10% < MCI < 90% | 79,99 | 67,58 | 55,35 |  | 80,00 | 79,33 | 65,85 |  | 80,04 | 94,16 | 85,22 |  | 80,18 | 33,56 |
| MCI > 90% | 9,97 | 1,56 | 1,13 |  | 9,99 | 6,93 | 2,85 |  | 9,98 | 0,18 | 0,29 |  | 9,91 | 0,06 |
| S < 10% | 10,00 | 1,56 | 1,13 |  | 9,97 | 6,93 | 2,85 |  | 9,98 | 0,18 | 0,29 |  | 9,91 | 0,19 |
| 10% < S < 90% | 80,02 | 67,59 | 55,41 |  | 80,09 | 79,35 | 65,87 |  | 80,11 | 94,16 | 85,22 |  | 80,18 | 44,38 |
| S > 90% | 9,97 | 30,85 | 43,45 |  | 9,95 | 13,72 | 31,28 |  | 9,91 | 5,66 | 14,49 |  | 9,91 | 55,43 |

**Table S2. Proportion of simple and complex mitochondria of COX-normal, intermediate and deficient fibres from all patients.**

*Table represent the 10th and 90th percentiles for volume, MCI and sphericity of mitochondria from COX-normal. Small and simple mitochondria with volume and MCI values inferior to the 10th percentile of mitochondria from COX-normal fibres. Large and complex mitochondria are those with volume and MCI values superior to the 90th percentile of mitochondria from COX-normal. Spherical mitochondria are those with values superior to the 90th percentile of mitochondria from COX-normal*.
